# Combinatorial transcription factor profiles predict mature and functional human islet α and β cells

**DOI:** 10.1101/2021.02.23.432522

**Authors:** Shristi Shrestha, Diane C. Saunders, John T. Walker, Joan Camunas-Soler, Xiao-Qing Dai, Rachana Haliyur, Radhika Aramandla, Greg Poffenberger, Nripesh Prasad, Rita Bottino, Roland Stein, Jean-Philippe Cartailler, Stephen C. J. Parker, Patrick E. MacDonald, Shawn E. Levy, Alvin C. Powers, Marcela Brissova

## Abstract

Islet-enriched transcription factors (TFs) exert broad control over cellular processes in pancreatic α and β cells and changes in their expression are associated with developmental state and diabetes. However, the implications of heterogeneity in TF expression across islet cell populations are not well understood. To define this TF heterogeneity and its consequences for cellular function, we profiled >40,000 cells from normal human islets by scRNA-seq and stratified α and β cells based on combinatorial TF expression. Subpopulations of islet cells co-expressing *ARX/MAFB* (α cells) and *MAFA*/*MAFB* (β cells) exhibited greater expression of key genes related to glucose sensing and hormone secretion relative to subpopulations expressing only one or neither TF. Moreover, all subpopulations were identified in native pancreatic tissue from multiple donors. By Patch-seq, *MAFA*/*MAFB* co-expressing β cells showed enhanced electrophysiological activity. Thus, these results indicate combinatorial TF expression in islet α and β cells predicts highly functional, mature subpopulations.

## INTRODUCTION

Pancreatic islets are cell clusters dispersed throughout the pancreas, composed primarily of endocrine cells that coordinate glucose homeostasis. Islet β cells secrete insulin which acts to lower blood glucose and α cells secrete glucagon which acts to raise blood glucose. In addition to α and β cells, cooperative interaction of less prevalent endocrine cells (δ, γ, and ε) and non-endocrine cell populations in the islet microenvironment, including endothelial cells, macrophages, pericytes (stellate cells), nerve fibers, and immune cells, provide additional signals to modulate islet function^1^. Islet α and β cells are characterized by the precise expression of transcriptional and signaling machinery that allows sensing and integration of glucose, nutrient, and neurohormonal signals and proportional response with regulated hormone secretion. Importantly, pancreatic islet dysfunction through impaired insulin and/or glucagon secretion is a hallmark of most forms of diabetes^2–5^. Thus, identifying key factors and molecular pathways governing α and β cell identity and function is crucial to understanding, treating, and preventing diabetes.

One set of important molecules governing α and β cell identity and function are islet-enriched transcription factors (TFs) that have been shown to have important roles in both islet development as well as in the maintenance of the islet cell phenotype, particularly in mouse and islet-like cells derived from human stem cells^6–9^. Importantly, several islet-enriched TFs have species differences between human and mouse, highlighting the need to closely investigate transcription factors in human systems^10, 11^. For example, members of the Maf transcription factor family show differences in cell type distribution and timing of expression^12, 13^. Such TFs interact in complexes and networks to exert broad control over cellular processes, making them foundational regulators of cell states. In fact, in addition to their coordinated role in islet cell development, loss or misexpression of key TFs has been highlighted in numerous forms of diabetes^14–17^.

Importantly, with advances in scientific methodologies, it has been increasingly recognized that islet cells are heterogeneous. This is particularly apparent in β cells, where recent work has highlighted human β cell heterogeneity in function^18^, cell surface protein expression^19, 20^, and transcriptomic profile^21, 22^. In contrast, heterogeneity within human α cells has been much less studied. Given the central role for islet-enriched TFs in regulating cell states, potential heterogeneity in these TFs may represent distinct cellular states with broad implications for human islet biology and diabetes.

RNA sequencing (RNA-seq) has been an essential technology to broadly characterize islet gene expression in an unbiased manner. Hallmark gene transcripts and gene pathways have been analyzed both at the whole islet level^23, 24^ and in a cell type-specific manner using fluorescence-activated cell sorting (FACS) with either cell surface markers on live cells or intracellular proteins in fixed and permeabilized cells to obtain purified α and β subpopulations^25–27^. However, these approaches provide limited ability to assess heterogeneity within a given cell type. To address this, single cell RNA-seq (scRNA-seq) is an exciting and evolving technology that can be used to understand cell type heterogeneity and has begun to be applied to human islets^18, 28–32^. While the magnitude of high-resolution data from these studies is exciting, there are also important technical challenges inherent to the small scale of input material^33, 34^, highlighting the importance of a robust comparison between bulk and scRNA-seq. Further, it remains unclear how α and β cells identified by protein-based methods (*e.g.*, FACS) compare to cells characterized by the clustering approach applied in scRNA-seq that arranges cells by transcriptional similarity.

To investigate how heterogeneity of islet-enriched TFs in human islets relates to islet function, we focused on three transcription factors, namely *ARX*, *MAFB*, and *MAFA*. ARX and MAFB are enriched in islet α cells, as are MAFA and MAFB in β cells, and all three play important roles in islet cell development and disease as suggested by existing bulk RNA-seq datasets^10, 17, 26, 27^.

Since our goal was to understand single cell heterogeneity, we translated findings from a bulk context to a single cell context by systematically analyzing the same islet preparation by both approaches to establish congruency between bulk and scRNA-seq methods. Finally, we generated a scRNA-seq dataset of over 40,000 islet cells from adult donors, which includes endocrine, immune, and endothelial cell populations, that is accessible through a user-friendly web portal. This dataset provided sufficient cell numbers to classify α and β cells into subgroups based on combinatorial *ARX*/*MAFB* and *MAFA*/*MAFB* expression, respectively, and allowed us to identify key correlates to α and β cell function. We further validated the existence of these cell populations within human pancreatic tissue *in situ* and linked *MAFA*/*MAFB* transcriptional heterogeneity of human β cells to their electrophysiological properties.

## RESULTS

### Transcriptional and immunohistochemical profiling of human α and β cells suggests a role for key transcription factors ARX, MAFA, and MAFB in islet cell development and disease

Both *in vivo* and *in vitro* studies have helped identify TFs with cell-specific expression patterns in islets. In α cells, Aristaless Related Homeobox (ARX) factor is essential for α cell differentiation and function, a finding which has been confirmed in human α cells^8, 35–37^. Indeed, *ARX* transcripts are heavily enriched in α cells (**Figures 1A-B and S1A-B**)^12, 38, 39^. Of note, α cells from donors with type 1 diabetes (T1D) show decreased *ARX* expression compared to α cells from nondiabetic donors (ND) (**Figure 1C**), indicating that this factor may contribute to impaired glucagon secretion observed in T1D^17, 18, 40^.

**Figure 1.**
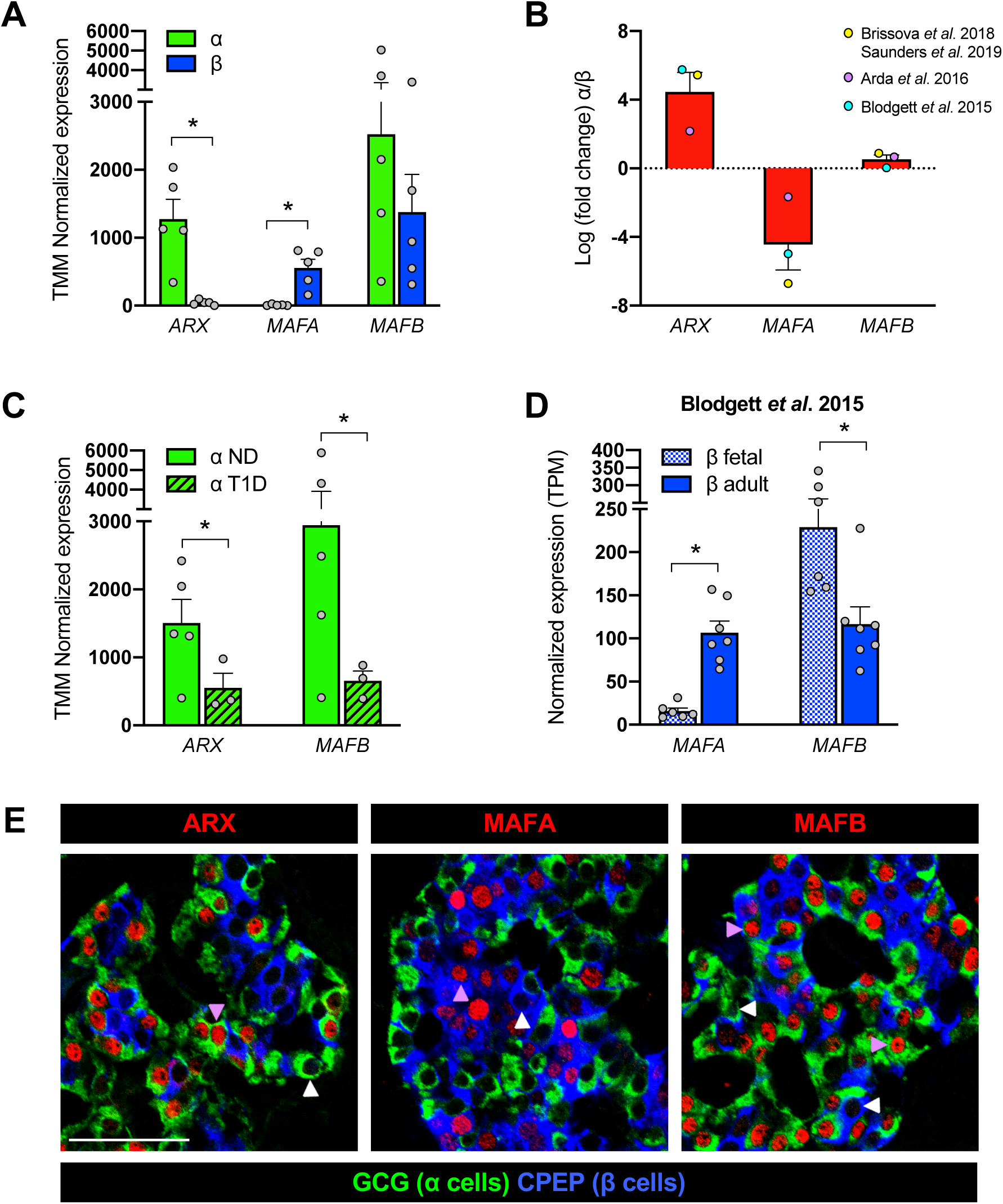
Bulk RNA-sequencing and immunohistochemistry data highlight unique expression patterns of transcription factors ARX, MAFA, and MAFB in human α and β cells. (A–D) Normalized expression values (A, C-D) and fold change (B) of *ARX*, *MAFA*, and *MAFB* in previously published bulk RNA-sequencing (RNA-seq) datasets from α cells (green) and β cells (blue). Data in A is from Brissova *et al*. 2018^17^ and Saunders *et al*. 2019^27^ (n=5 donors); additional datasets Arda *et al*. 2016^10^ (n=5 donors) and Blodgett *et al*. 2015^26^ (n=7 donors) are included in panel B. See also Figure S1A-B. (C) Expression of *ARX* and *MAFB* is decreased (*ARX* fold change: -2.7; *MAFB*: -3.4) in α cells from donors with type 1 diabetes (T1D) compared to nondiabetic (ND) donors^17^. (D) Expression of *MAFA* is increased (fold change: 7.1) in adult β cells compared to fetal β cells, while *MAFB* is decreased (fold change: - 2.0)^26^. All bar graphs show mean + SEM; symbols represent individual donors (panels A, C-D) or average value per dataset (B). Asterisks indicate significant (adjusted p-value <0.05) fold change of α vs. β in panels A and B, T1D vs. ND in C, and adult vs. fetal in D. (E) Immunohistochemical staining of pancreatic sections from a nondiabetic adult (55 years, **Table S4**), showing specificity of ARX, MAFA, and MAFB (red) in α cells (GCG; green) and β cells (CPEP; blue). Arrowheads indicate cells negative (white) or positive (purple) for transcription factors; scale bar, 50 μm. See also Figure S1C-D.

MAFA is a *bona fide* β cell factor exerting direct control over both insulin expression as well as key components of glucose-stimulated insulin secretion, and it is expressed relatively late in β cell development, making it a commonly used marker of fully mature β cells^41–43^. MAFA is thought to play a broadly similar role in adult mouse and human β cells, and existing RNA-seq datasets underscore its β cell specificity (**Figures 1A-B** and **S1A-B**). MAFA is clearly present in adult β cells but its expression actually does not peak until several years after birth, as illustrated by previous histological studies^12^ and transcriptomic profiles of β cells from fetal versus adult donors (**Figure 1D**)^26^. These data temporally correlate increased MAFA levels with the acquisition of increased glucose sensitivity^44–46^, suggesting that MAFA plays a role in β cell maturation and function.

In contrast to ARX and MAFA, MAFB is expressed by both α and β cells (**Figures 1A-B** and **S1A-B**) and shows significant species differences: it is retained in human β cells during adulthood, while in rodents it becomes restricted to α cells in the early postnatal period^11^. Of note, the MAF factors are thought to be capable of forming both homo- and heterodimers^47^, providing an opportunity for synergy between MAFA and MAFB in β cells. In α cells, MAFB is known to directly bind to the *GCG* promoter to regulate glucagon expression^32^, rendering it an important regulator of α cell function. Like *ARX*, *MAFB* is reduced in α cells from donors with T1D (**Figure 1C**).

The unique and dynamic expression patterns of *ARX*, *MAFA*, and *MAFB* demonstrated by bulk RNA-seq (**Figures 1A-1D** and **S1A-B**) suggest that these TFs are linked to key aspects of α and β cell function. However, our analysis of their special distribution in adult human pancreatic tissue revealed that not all α or β cells in a given islet express them (**Figures 1E** and **S1C-D**). Thus, to further understand the role of these TFs, we sought to determine the cell-to-cell variability that cannot be discerned from a pooled cell population profiled by bulk RNA-seq.

Given the known importance of TFs in regulating cellular processes, we hypothesized that TF heterogeneity at the single cell level could define α or β cell subtypes with different functional properties.

### Gene expression profiles obtained by scRNA-seq are largely concordant with those obtained by bulk RNA-seq

To translate gene expression findings from a FACS-sorted bulk context to a single cell context, we systematically analyzed FACS-purified α and β cells from a healthy 39-year-old donor by the two approaches in parallel (**Figures 2A**, **S2A**; **Table S1**). Performing all analyses on the same donor allowed us to avoid donor-to-donor variability. Approximately 10,000 cells for each cell type were pooled to generate bulk RNA-seq libraries (“FACS-Bulk-α” and “FACS-Bulk-β”) while ≥10,000 α cells and β cells went to scRNA-seq to capture 6,371 and 1,190 single cells after quality control (“FACS-SC-α” and “FACS-SC-β”), respectively. We compared expression of all genes between bulk RNA-seq vs. pooled single cells (pseudo-bulk) from scRNA-seq and as expected, found that expression tended to be higher in the bulk compared to single cell samples. To broadly understand gene expression differences between the two approaches, we compared genes detected above 1 log2TPM (transcript per million) for both (**Figures 2B, 2C**). Of the genes in the FACS-SC-α or FACS-SC-β groups, approximately 97% were also in FACS-Bulk-α or FACS-Bulk-β, respectively. In contrast, just 8.5% of genes in FACS-Bulk-α and 6.5% of genes in FACS-Bulk-β were also in the respective single cell dataset.

**Figure 2.**
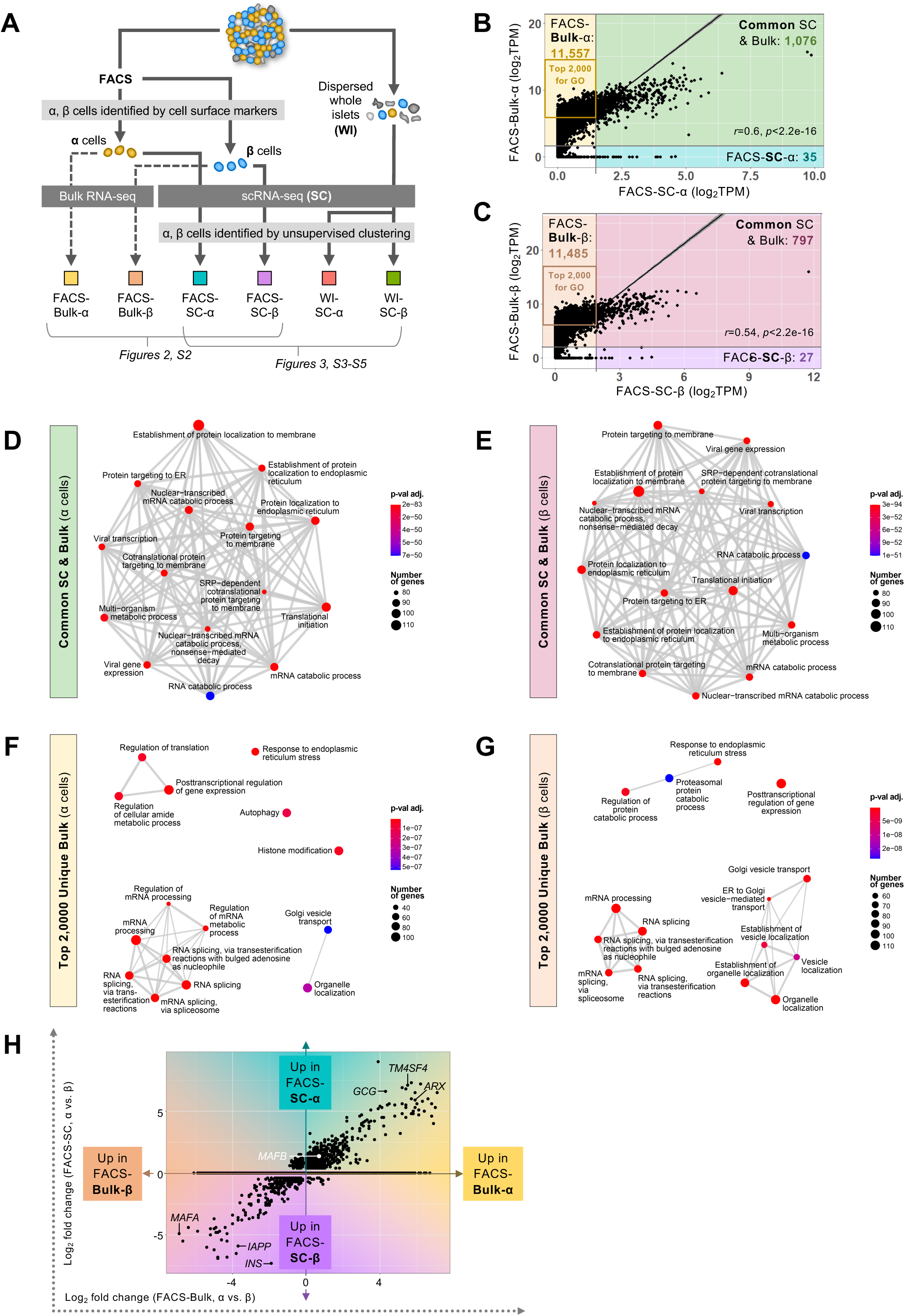
Similarities and differences in gene capture between single cell and bulk RNA-seq. **(A)** Schematic showing the comparison of sorted human α and β cells profiled by bulk (FACS-Bulk) and single cell (FACS-SC) RNA-sequencing (n=1, 39y donor; data in Figures 2 and S2), as well as single α and β cells identified by cell surface markers (FACS-SC) compared to those from dispersed whole islets (WI-SC) identified by unsupervised clustering. (n=2, 14y and 39y donors; data in Figures 3 and S3**-S5**). **(B–C)** Average expression (TPM) was taken across 7,269 α cells and 2,511 β cells from scRNA-seq and compared with TPM normalized expression of bulk RNA-seq (10,000 cells/each) of corresponding populations. Only genes above log2TPM=1 in both populations were considered to assess gene detection; *r* is Pearson’s coefficient and *p* is significance from t-test statistic. **(D–G)** Gene ontology analysis was performed on genes common between scRNA-seq and bulk RNA-seq (**D**, α cells; **E**, β cells), as well as on the 2,000 most highly expressed genes unique to bulk RNA-seq (**F**, α cells; **G**, β cells) using “enrichDAVID” function of the R package clusterProfiler 3.14.3^64^. Colored labels show data input and correspond to shaded regions of panels **B**-**C**. **(H)** Comparison of individual genes differentially expressed between α and β cells, with log2 fold change from scRNA-seq plotted on y-axis and bulk RNA-seq on x-axis.

To characterize molecular pathways captured by scRNA-seq and determine what additional information may be captured uniquely in bulk RNA-seq, we analyzed ontologies from the genes that were common between bulk and single cell samples (green in **Figure 2B** and pink in **Figure 2C**) as well as the 2,000 most highly expressed genes uniquely captured by bulk RNA-seq (yellow box in **Figure 1B** and orange box in **Figure 1C**). Visualization of these ontologies in an enrichment map highlighted that the common genes covered a broad and comprehensive range of biological processes (**Figures 2D-E**). The additional processes represented by highly expressed genes unique to bulk RNA-seq were, by contrast, less enriched (**Figures 2F-2G**).

Indeed, visualization of the top 30 most significantly enriched processes highlighted a higher degree of enrichment in the shared common and bulk group (**Figures S2B**-**C**). Biological processes associated with hormone secretory function, such as regulation of insulin secretion or ER to Golgi vesicle-mediated transport, were represented in all data sets (**Figure S2D-E**).

As bulk RNA-seq and scRNA-seq involve different chemistries that may bias direct comparisons of gene expression levels, we next assessed relative differences by looking at differential expression between α and β cells profiled by each approach (Bulk-α vs. Bulk-β compared to pooled data from SC-α vs. SC-β). Genes differentially expressed in both datasets were highly correlated (*r*=0.91, *p*<2.2e-16) and showed the expected enrichment of β cell-specific genes (*e.g.*, *INS, IAPP*) as well as α cell-specific genes (*GCG, TM4SF4*) (**Figure 2H**). Importantly, there were very few differentially expressed genes that were regulated in opposite directions (top left and bottom right quadrants in **Figure 2H**), suggesting that trends in gene expression are consistent between the two methods. Taken together, these data indicate that although bulk RNA-seq captures a greater breadth of genes, scRNA-seq analysis captures a similarly broad and comprehensive set of pathways and processes that are specific to α and β cell biology.

### Gene expression profiles of α and β cells identified by unsupervised clustering are consistent with profiles of α and β cells resolved by cell surface markers

We next asked whether unsupervised clustering (identification of cells post-sequencing) yielded similar gene expression profiles to those cells identified and obtained by sorting with previously characterized cell surface markers^17, 48, 49^. Dispersed islet cells from two healthy donors were profiled by scRNA-seq after either purification into α and β cell populations by FACS (“FACS-SC-α” and “FACS-SC-β”) or directly from hand-picked whole islets dispersed without any additional purification (“WI-SC-α” and “WI-SC-β”; **Figure 2A**; **Table S1**). A total of 27,614 cells across the four groups passed quality control (see **Methods**). Cells from dispersed whole islets (**Figure S3A**), FACS-α (**Figure S3B**), and FACS-β (**Figure S3C**) were analyzed by graph-based unsupervised clustering applying Louvain algorithm^50, 51^ and visualized using Uniform Manifold Approximation and Projection (UMAP)^52^ and α and β cells were annotated with markers (**Table S2**) overlayed to unsupervised clusters. Total gene capture was similar across all four cell populations analyzed (median 2,068 genes per cell; **Figure S3D**). Principal Component Analysis (PCA) of all four samples indicated that overall variance is governed by cell type differences rather than the approach used to stratify α and β cells (cell sorting vs. unsupervised clustering) (**Figure 3A**; individual donors shown in **Figure S4A**). This was also apparent by looking at single cell expression level of the genes with greatest influence on principal component 1 (**Figure 3B**; individual donors shown in **Figure S4B**). These genes included both known α and β cell markers (*e.g.*, *GCG*, *SLC7A2*, *GC*, *INS*, *PCSK1*) as well markers that have been previously identified but not extensively studied in islets (*e.g.*, *RGS4*, *FXYD3*, *FAP*, *MEG3*, *HADH*, *SAMD11*).

**Figure 3.**
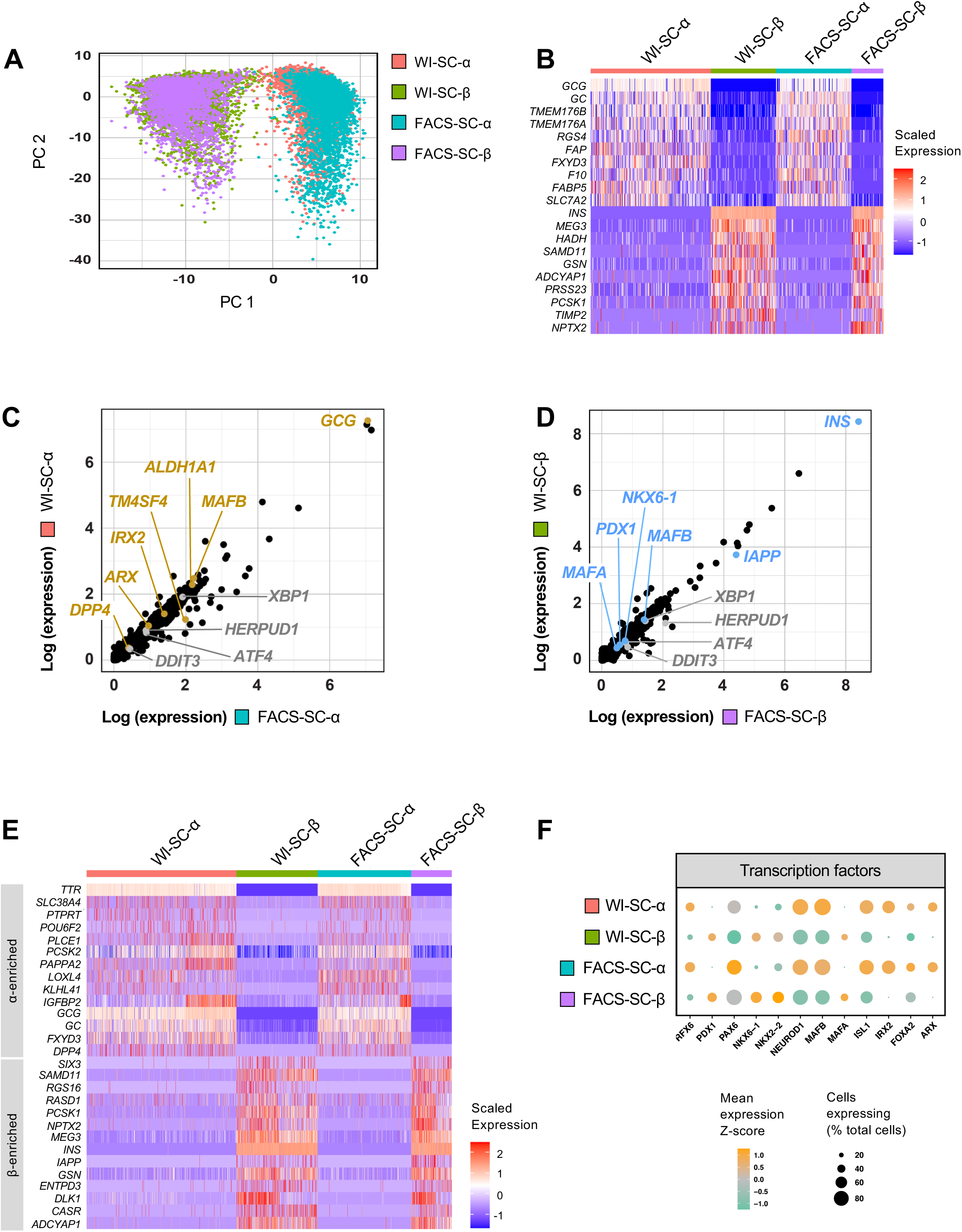
Gene expression of α and β cells by scRNA-seq is similar between cells identified with cell surface markers and those identified by unsupervised clustering. **(A)** Principal component analysis (PCA) shows clustering of sorted α and β cells identified by cell surface marker expression (FACS-SC) and those derived from dispersed whole islets and identified by unsupervised clustering (WI-SC). See also Figure 2A. **(B)** Heatmap depicts expression of those genes contributing to variability in PCA. **(C–D)** Comparison of average log expression of genes across cells identified by unsupervised clustering or cell surface markers for α (**C**) and β cells (**D**). Genes highlighted are α cell-enriched (yellow), β cell-enriched (blue), or selected markers of cell stress (grey). **(E)** Heatmap showing variable expression of known α and β cell-enriched markers within and between each sample. **(F)** Relative expression of transcription factors across samples; dot size indicates the percentage of cells with detectable transcripts and color indicates gene’s mean expression z-score.

Gene expression profiles of FACS-α and FACS-β samples showed strong linear correlation (Pearson’s correlation, *r*=0.99, p<2.2e-16) with WI-α and WI-β samples, respectively (**Figure 3C-D**; individual donors shown in **Figure S5A-B**). Of note, α and β cell-enriched genes, including transcription factors, and stress markers were all expressed, on average, at similar levels between WI and FACS samples in both islet preparations studied. To appreciate cell heterogeneity, we visualized canonical α and β cell markers (**Figure 3E**; individual donors shown in **Figure S5C**) which highlighted that both approaches demonstrated similar variability within these key genes. Finally, we visualized key islet-enriched transcription factors in a dot plot by the number of cells expressing the factor and the average normalized expression level, and we found consistent results between the two methods (**Figure 3F**). Thus, these results indicate that the cell sorting step does not alter the transcriptional profile of the FACS-purified α and β cells. They further suggest that post hoc identification of cell types by unsupervised clustering based on transcriptional profile is consistent to the well characterized approaches of cellular identification using antibodies to cell surface proteins by FACS. Thus, both approaches are likely identifying the same cell populations and would allow investigation of TF heterogeneity.

### scRNA-seq reveals heterogenous transcription factor expression in α and β cells

One major advantage of scRNA-seq is its ability to dissect heterogeneous cell composition within and across cell types. However, because some subpopulations are relatively rare, robust datasets are required to sufficiently characterize these populations. In this study, we obtained 44,953 high-quality single cell transcriptomes of hand-picked islets from n=5 healthy donors with robust dynamic insulin and glucagon secretion profiles characterized by perifusion to ensure healthy and functional cells were being assessed (**Table S1** and **Figure S6A**). Graph-based unsupervised clustering^50^ reliably detected major endocrine cell types (α, β, δ) and also acinar, ductal, stellate, endothelial, and immune cells (**Figure 4A**). Clusters were annotated to identify cell types, including rare populations such as γ and ε, using markers listed in **Table S2** and identified cell types were represented in each donor (**Figure S6B**). Cell populations were confirmed by the specific expression of additional known identity markers (**Figure 4B**). Within cell types, the expected clustering by individual donor (**Figure S6C**) is apparent. To facilitate the exploration of this robust single cell dataset, we created an online application that allows one to browse single cell gene expression by both the cell type and donor (**Figure S6D**).

**Figure 4.**
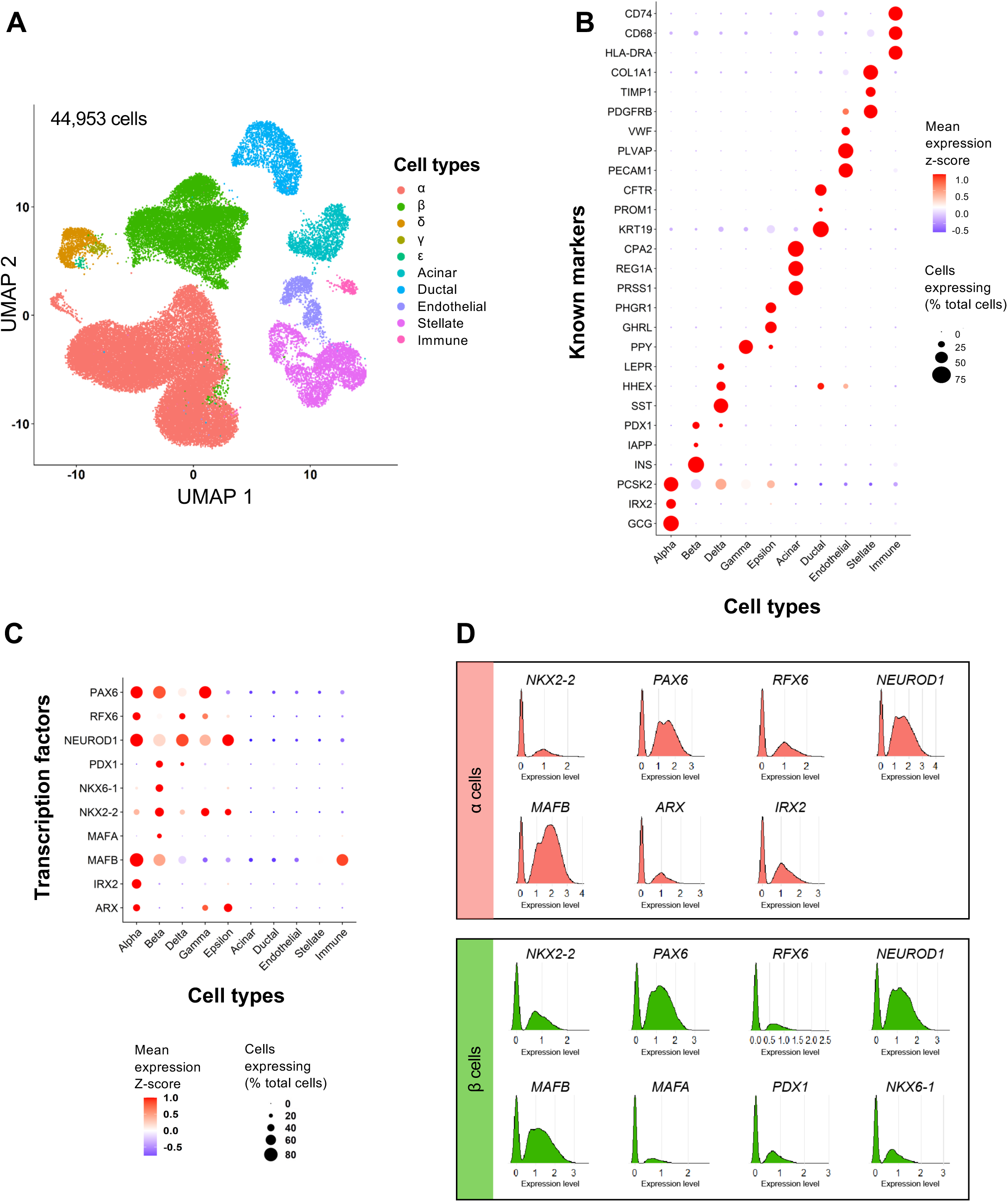
Transcription factor expression in human pancreatic islets by scRNA-seq. **(A)** UMAP visualization of 44,953 pancreatic islet cells from n=5 islet preparations, identified by unsupervised clustering; cell populations include β (24%), α (54%), δ (2.5%), ε (0.08%), acinar (3.3%), ductal (4.7%), endothelial (2.2%), stellate (7.7%), and immune cells (0.5%). Cell clusters were annotated using known gene markers (**Table S2**). γ and ε cells could not be resolved from the δ cell cluster; thus, these populations were manually selected using the “CellSelector” function to identify cells positive for *PPY* and *GHRL*, respectively. Libraries were sequenced at ∼80,000 reads/cell yielding a median of 2,365 genes per cell. **(B)** Dot plot showing relative expression of cell type markers to validate cell type annotation post-unsupervised clustering. **(C)** Dot plot showing relative expression of transcription factors across all cell types. In panels **B**-**C**, dot size indicates the percentage of cells with detectable transcripts; color indicates gene’s mean expression z-score. **(D)** Natural log expression level of common transcription factors expressed in α and β cells.

To investigate the cell-specific signatures of human α and β cells, we analyzed expression patterns of canonical islet-enriched TFs. *PAX6*, *RFX6*, *NEUROD1*, and *NKX2-2* were expressed in all endocrine cell types, whereas *PDX1*, *NKX6-1*, and *MAFA* were enriched in β cells, *IRX2* was specifically expressed in α cells, and *ARX* was expressed in α, γ, and ε cells, consistent with previous single cell studies^28, 29, 50^ (**Figure 4C**). *PAX6*, *NEUROD1* and *MAFB* were among the most prevalent endocrine factors, expressed in >75% of both α and β cells (**Figure 4C**). Of particular interest, *MAFB* – known in humans to be expressed in both α and β cells – is also enriched in the immune cell population, which had been overlooked in previous studies due to low abundance of immune cells in isolated islets. Interestingly, we noticed that each of these key TFs had a bimodal distribution, meaning there was a clear subpopulation of cells without detectable expression of each factor (**Figure 4D**), consistent with our observations for MAFA, MAFB and ARX in pancreas tissue (**Figure 1E**). Given the crucial role islet-enriched TFs play in islet cell identity and function, particularly when acting in TF regulatory networks, we thus hypothesized that combinations of key TFs would identify important islet cell subtypes.

### Heterogeneity of *ARX* and *MAFB* expression in α cells by scRNA-seq predicts expression of key α cell functional genes

Since both ARX and MAFB are downregulated in α cells from donors with T1D^17^, we tested the hypothesis that these factors cooperatively regulate α cell function. We first confirmed heterogeneous *ARX* and *MAFB* expression in α cells from all five donors (**Figure 5A**). Of 24,248 total α cells, we identified populations of α cells without *ARX* or *MAFB* expression (“None;” 10%), populations expressing only *ARX* or only *MAFB* (4% and 48%, respectively), and a population co-expressing both *ARX* and *MAFB* (“Both;” 38%) that were relatively stable across all five donors (**Figure 5B**). For these four populations, we investigated expression of other islet-enriched TFs, α cell-enriched genes, and genes related to ion flux, glucose metabolism, vesicle trafficking, exocytosis and cell stress (**Figures 5C** and **S7**). Interestingly, we observed that numerous α cell-enriched TFs (*RFX6*, *PAX6*, *NEUROD1*, *ISL1*, *IRX2*) and genes related to nutrient sensing or glucagon secretion (*ACLY*, *PKM*, *GSTA4*, *GPX3*, *G6PC2*, *KCTD12*, *KCNK16*, *KCNJ6*, *ABCC8*) were elevated in α cells co-expressing *MAFB* and *ARX* compared to the other populations, while genes related to cell stress (*DDIT*, *ATF4*) were highest in the “None” group, suggesting that presence of both factors may support increased metabolic activity and glucagon secretory capacity. To confirm these findings, we analyzed three additional scRNA-seq datasets of human islets that utilized different single cell technologies^18, 28, 29^ and found the results to be consistent (**Figure S8A**).

**Figure 5.**
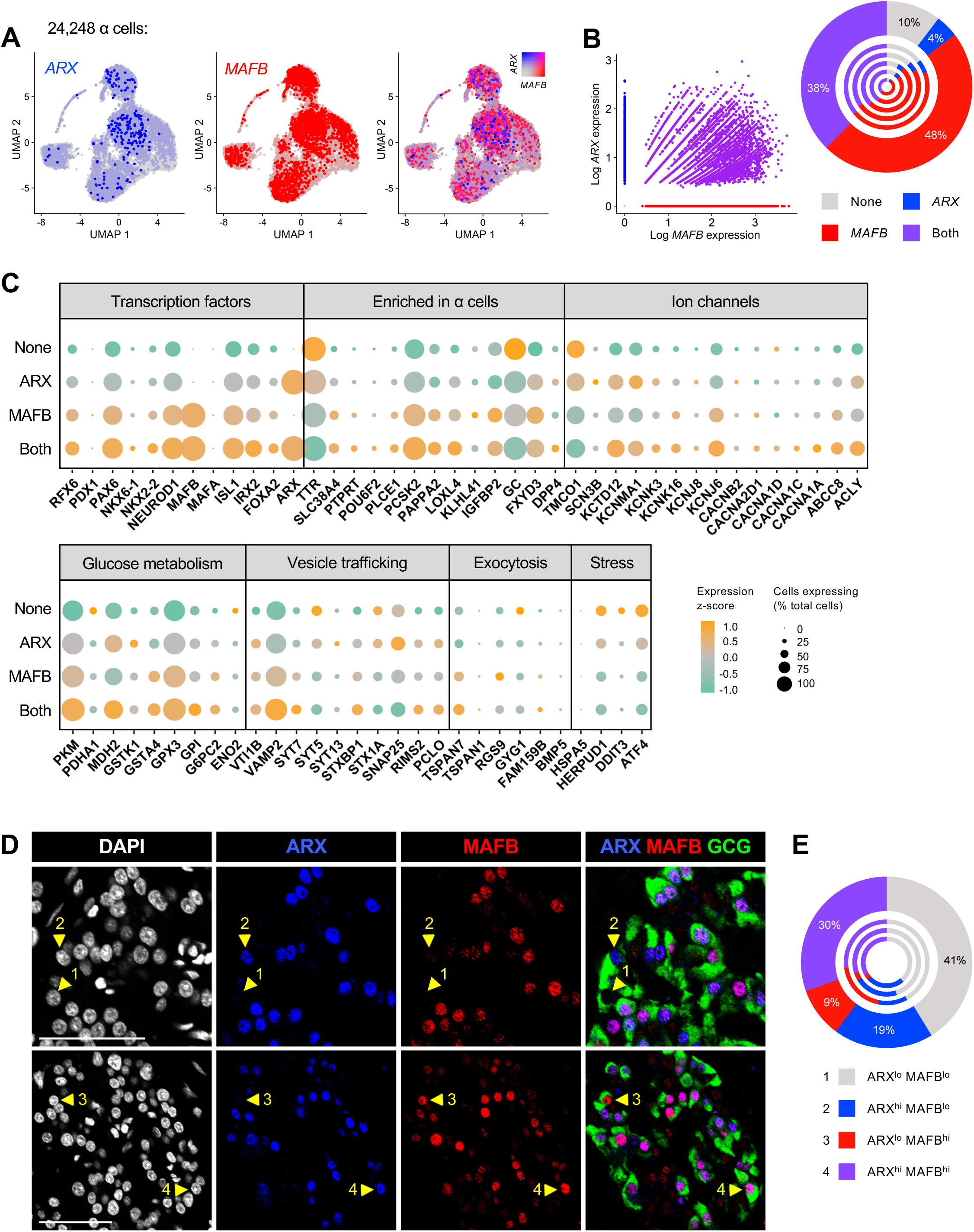
Heterogeneity of *ARX* and *MAFB* expression in α cells by scRNA-seq correlates with expression of key functional genes. **(A)** UMAP visualization of 24,248 α cells (n=5 donors) pseudocolored to show, from left to right, expression of *ARX* (blue); *MAFB* (red); and both *ARX* and *MAFB* with 0.5 color threshold scale. **(B)** Scatterplot on the left is depicting four distinct α cell populations based on *ARX* and *MAFB* expression: those expressing neither factor (10%), those expressing only *ARX* (4%) or only *MAFB* (48%), and those co-expressing *ARX* and *MAFB* (38%). Chart on the right shows cell populations by donor, with the outermost circle reflecting totals. **(C)** Dot plot showing the relative expression of selected genes related to α cell identity, ion flux, glucose metabolism, vesicle trafficking, exocytotic machinery, and cellular stress of the four α cell populations in panel **B**. Dot size indicates the percentage of α cells with detectable transcripts; color indicates the gene’s mean expression z-score. See **Figure S8** for comparison to other single cell studies. **(D)** Immunohistochemical staining of ARX (blue) and MAFB (red) in glucagon (GCG)-expressing α cells (green) of a nondiabetic adult (55 years, **Table S4**). Numbered arrowheads indicate the presence of 4 α populations: 1, ARX^lo^ MAFB^lo^; 2, ARX^hi^ MAFB^lo^; 3, ARX^lo^ MAFB^hi^; 4, ARX^hi^ MAFB^hi^. **(E)** Quantification of α cell populations shown in panel **D** (n= 2,369 α cells). Outermost circle represents composite count and inner circles represent α cells from each of n=3 donors (see also **Figure S7B**).

We next asked whether *ARX*/*MAFB* heterogeneity existed at the protein level given the known differences that exist between transcript and protein expression^53^. To assess this, we performed immunohistochemical analysis of ARX and MAFB on pancreatic tissue sections from nondiabetic donors (**Figures 5D** and **S8B**). Cells were classified by automated algorithm for “low” or “high” ARX and MAFB expression, setting an intensity threshold that remained consistent across all islets from a given tissue. By this measure, all four combinations of ARX/MAFB-expressing α cells were detected in each donor evaluated: ARX^lo^ MAFB^lo^ (41%), ARX^hi^ MAFB^lo^ (19%) ARX^lo^ MAFB^hi^ (9%) and ARX^hi^ MAFB^hi^ (30%) (**Figure 5E**). Taken together, our results indicate the presence of α-cell subpopulations classified according to unique and conjunctional expression of ARX and MAFB and suggest that combined expression of these two markers likely identifies highly functional and mature α cells.

### β cells co-expressing *MAFA* and *MAFB* exhibit characteristics of enhanced secretory function

Given the ability of MAFA and MAFB to heterodimerize^47^ and the unique expression changes during β cell maturation^10, 12, 26^, we hypothesized that MAFA and MAFB co-expression represents a unique subpopulation of human β cells. To test this, we resolved 11,034 β cells into subgroups that expressed only *MAFA* or only *MAFB* (4% and 52%, respectively), β cells that co-expressed both *MAFA* and *MAFB* (“Both;” 22%), and β cells with undetected expression of *MAFA* and *MAFB* (“None;” 21%) (**Figure 6A-B**). We assessed these groups for the same set of key cellular identity and functional genes described above for α cells, and we saw a general trend of increased expression of key functional genes with dual *MAFA* and *MAFB* expression (**Figures 6C** and **S9**). Specifically, numerous genes related to cell identity (*PDX1*, *PAX6*, *NEUROD1*, *ISL1*, *PCSK1*, *IAPP*), glucose metabolism (*ACLY*, *G6PC2*, *GPX3*), ion channels (*ABCC8*, *KCNJ6*), and exocytosis (*VAMP2*, *SYT7*, *PCLO*, *TSPAN7*, *RGS9*, *FAM159B*, *BMP5*) were all increased in MAFA and MAFB co-expressing cells compared to other subgroups. In contrast, stress genes (*HSPA5*, *HERPUD1*, *DDIT3*, *ATF4*) were either significantly reduced in the co-expression group or significantly elevated in the “None” group. These expression patterns indicate that presence of both factors may be crucial for increased metabolic activity and insulin secretion. Analysis of three independent single cell studies of human islets utilizing other platforms^18, 28, 29^ confirmed these results (**Figure S10A**). The presence of β cell MAFA/MAFB heterogeneity at the protein level (MAFA^lo^ MAFB^lo^, 46%; MAFA^hi^ MAFB^lo^, 8%; MAFA^lo^ MAFB^hi^, 29%; MAFA^hi^ MAFB^hi^, 16%) was validated by immunohistology in pancreatic sections, where cells representative of all four populations were identified in each of multiple non-diabetic donors (**Figures 6D-E** and **S10B**).

**Figure 6.**
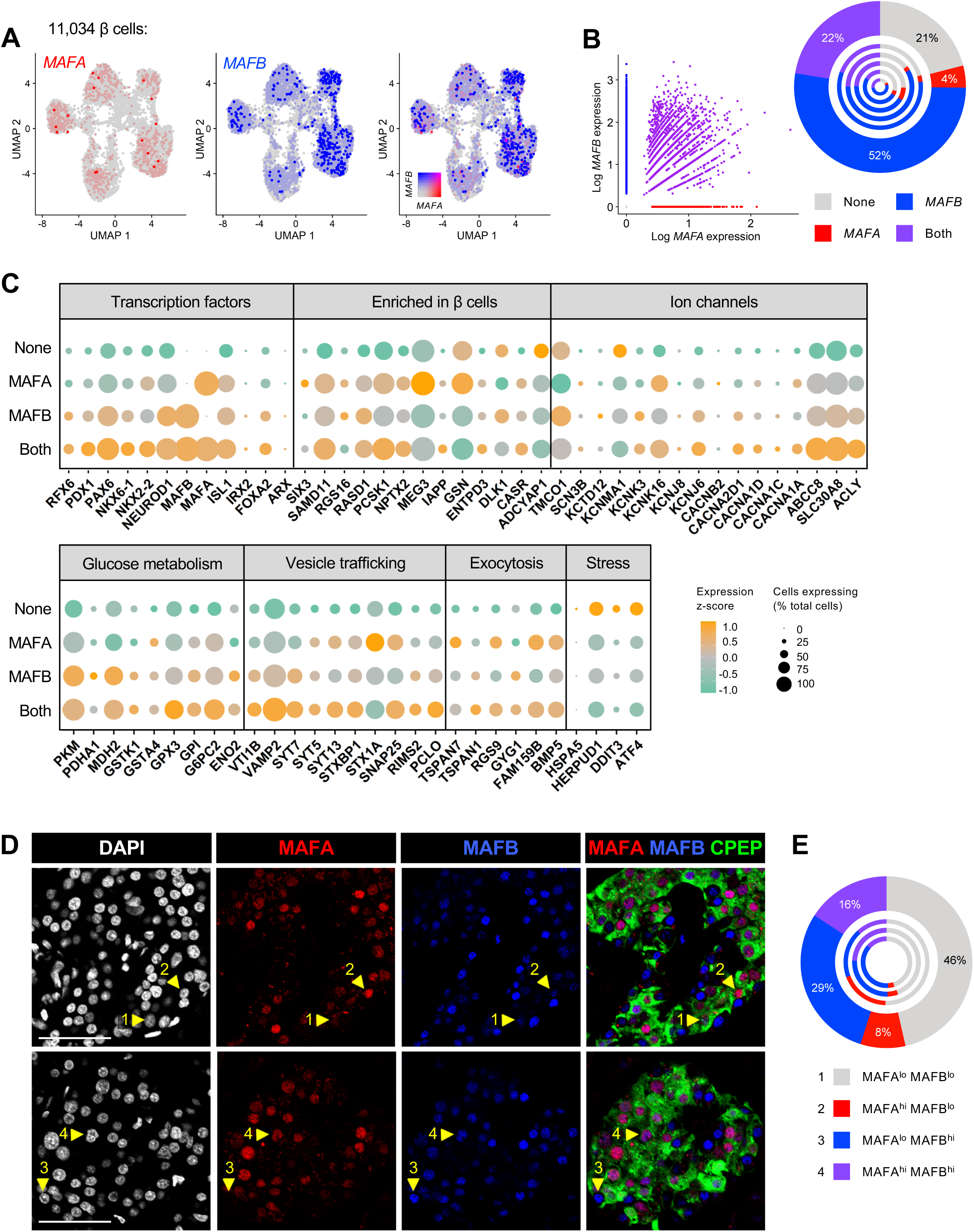
Heterogeneity of MAFA and MAFB expression in β cells by single cell RNA-seq correlates with expression of key genes involved in β cell function. **(A)** UMAP visualization of 11,034 β cells (n=5 donors), pseudocolored to show, from left to right, expression of *MAFA* (red); *MAFB* (blue); and both *MAFA* and *MAFB* with 0.5 color threshold scale. **(B)** Scatterplot on the left depicts four distinct β cell populations based on *MAFA* and *MAFB* expression: those expressing neither factor (22%), those expressing only *MAFA* (4%) or only *MAFB* (52%), and those co-expressing *MAFA* and *MAFB* (22%). Chart on the right shows cell populations by donor, with the outermost circle reflecting totals. **(C)** Dot plot showing the relative expression of selected genes related to β cell identity, ion flux, glucose metabolism, vesicle trafficking, exocytotic machinery, and cellular stress of the four β cell populations in panel **B**. Dot size indicates the percentage of β cells with detectable transcripts; color indicates the gene’s mean expression z-score. See **Figure S10** for comparison to other single cell studies. **(D)** Immunohistochemical staining of MAFA (red) and MAFB (blue) in C-peptide (CPEP)-expressing β cells (green) of a nondiabetic adult (55 years, **Table S4**). Numbered arrowheads indicate the presence of 4 populations: 1, MAFA^lo^ MAFB^lo^; 2, MAFA^hi^ MAFB^lo^; 3, MAFA^lo^ MAFB^hi^; 4, MAFA^hi^ MAFB^hi^. **(E)** Quantification of β cell populations shown in **D** (n= 2,566 β cells). Outermost circle represents composite count and inner circles represent β cells from each of n=3 donors (see also **Figure S8B**).

To determine whether the β cell subpopulation co-expressing *MAFA* and *MAFB*, enriched for numerous genes related to metabolism and hormone secretion, had functionally relevant consequences compared to other β cells, we utilized human Patch-seq data from Camunas et al.^18^. Transcriptomes from 194 β cells within this dataset (**Figure 7A**) showed high similarity with our larger dataset of 11,034 β cells (**Figures 6C** and **S10A**). In addition to producing an mRNA profile, the Patch-seq approach captures an electrophysiological profile of each cell, generating linked data on cell size, exocytosis, and ion channel currents. In agreement with transcriptome data, β cells that co-expressed both *MAFA* and *MAFB* showed increased electrophysiologic activity across several parameters including early exocytosis, early and late Ca^2+^ current, and late Ca^2+^ conductance when compared to cells that expressed *MAFA* only, *MAFB* only, or neither factor (**Figure 7B**). Of note, *MAFA*/*MAFB* co-expressing β cells are comparable in size to those expressing only one or neither factor, suggesting that neither the transcriptomic data nor the elevated electrophysiologic activity can be attributed to larger cells expressing more genes (**Figure 7B**). Thus, these data provide strong support that heterogeneous populations of β cells on the basis of combinatorial MAFA/MAFB expression exist and that co-expression of both factors marks β cells with elevated function.

**Figure 7.**
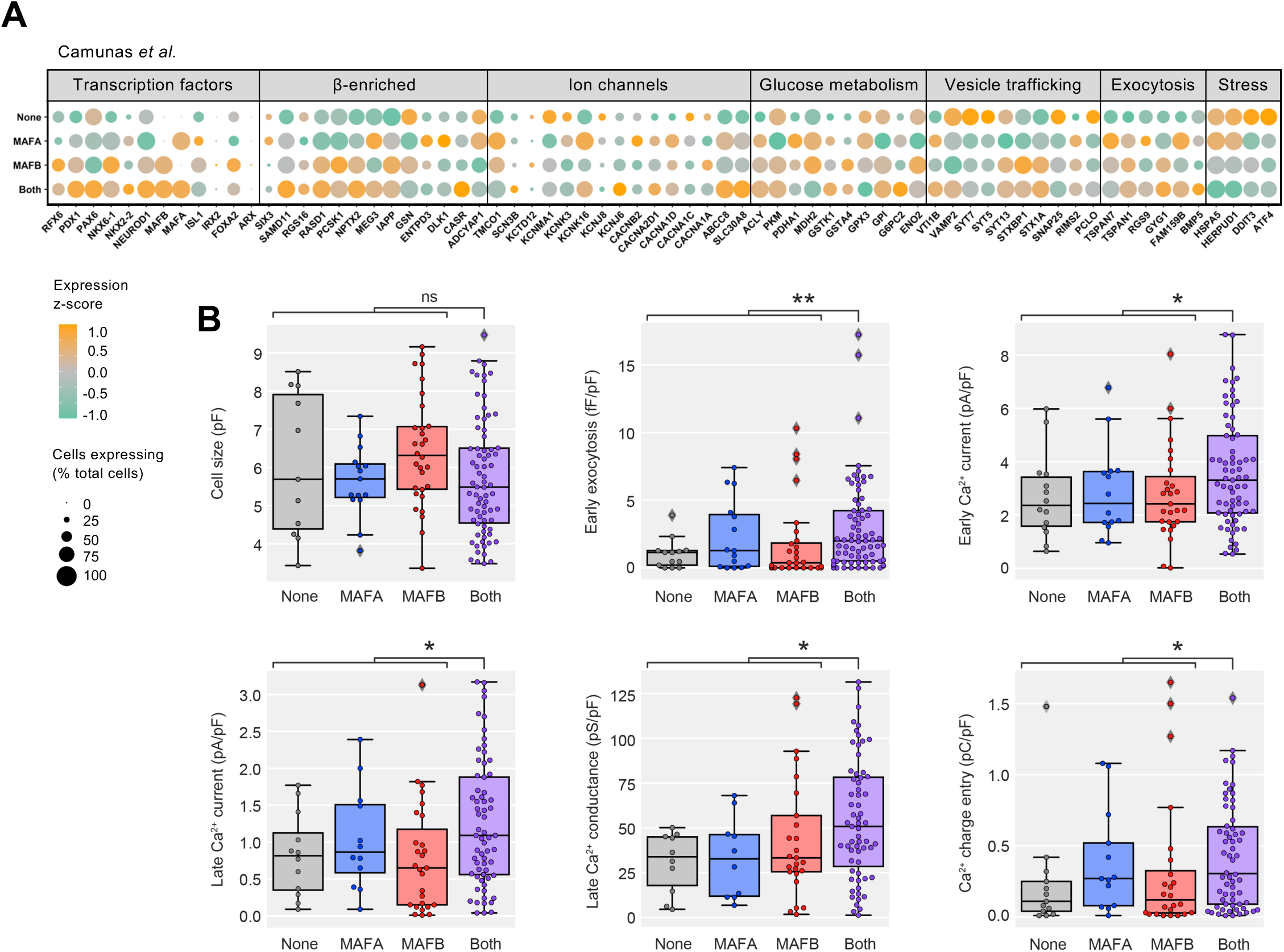
Beta cells co-expressing *MAFA* and *MAFB* have enhanced electrophysiolgic activity compared to β cells expressing one or neither factor. **(A)** Dot plot showing the relative expression of selected genes in β cells expressing neither *MAFA* nor *MAFB*, those expressing only *MAFA* or only *MAFB*, and those co-expressing *MAFA* and *MAFB*, based on data from Camunas et al. 2020^18^. Dot size indicates the percentage of cells with detectable transcripts; color indicates gene’s mean expression z-score. **(B)** Electrophysiological function in *MAFA*- and *MAFB*-expressing β cell subpopulations. Significantly higher Ca^2+^ currents and exocytosis are observed for β cells expressing both *MAFA* and *MAFB* with similar cell size across all subpopulations. Mann-Whitney test adjusted for multiple hypothesis testing with Benjamini-Hochberg (BH) procedure; *, p < 0.05; **, p < 0.01.

## DISCUSSION

By transcriptional profiling and assessment of protein expression at the single cell level, we found that several key islet-enriched TFs important for α and β cell maturity and function had a heterogenous expression pattern within normal human islet cells. To unravel the functional consequences of this heterogeneity in TF expression, we systematically analyzed the same islet preparation by bulk and scRNA-seq approaches and established congruency between the two methods. Capitalizing on our large scRNA-seq dataset, we stratified α and β cells based on differential or combined expression of key TFs (*ARX*/*MAFB* in α cells; *MAFA*/*MAFB* in β cells) that are known to act cooperatively. We found that co-expression of these TF combinatorial pairs predicted greater expression of genes related to glucose metabolism, ion flux, and hormone secretion, including both known α and β cell functional markers and those not extensively studied in islets. Importantly, we identified subpopulations with TF heterogeneity at the protein level by spatial analysis of normal human tissue and demonstrated, using Patch-seq, greater electrophysiological activity in *MAFA* and *MAFB* co-expressing β cells. These results suggest that combinatorial expression of key islet TFs defines highly functional and mature α and β cells.

Bulk RNA-seq and scRNA-seq have provided immense knowledge of the human islet transcriptional landscape, but each technology has strengths and drawbacks. Despite the prevalence of both approaches, this study is, to our knowledge, the first to report direct comparisons of bulk RNA-seq on FACS-purified human α and β cells and scRNA-seq on FACS-purified or dispersed cells from the same islet preparation. We highlight that while sensitivity to low expression genes is reduced in scRNA-seq, the detected genes cover a broad range of biological pathways that allow reconstructing gene ontology enrichment maps obtained from bulk RNA-seq (**Figure 2D-E**). Further, α and β cells show very similar expression profiles regardless of cell type identification method, with neither clustering via transcriptional similarity nor presence of characterized cell surface proteins showing an apparent bias. This indicates that enrichment methods using cell-surface markers are an appropriate method to investigate subpopulation of islet cell types.

Lower gene expression in scRNA-seq compared to bulk was an expected finding given that bulk RNA-seq generates reads from nearly the entire length of a gene, while the 10x platform, used in this study, does so only from the 3’ end. Single cell technologies that capture full length transcripts (*e.g.*, Smart-Seq2) may fare better in direct comparison of gene expression levels, though this hasn’t been investigated in islets^54^. Indeed, the smaller working range and lower signal-to-noise ratio is reflected in our scRNA-seq data. Despite this, transcripts above a TPM=1 threshold in both datasets converged linearly and were involved in a broad range of similar biological processes, emphasizing the high fidelity of both methods to assess islet cell biology (**Figure 2B-G**). To mitigate the differential scale, we also compared the relative transcript abundance in the form of α versus β cell enrichment (**Figure 2H**). Again, scRNA-seq was not as sensitive to changes across all transcripts but those that were detected exhibited very high correlation.

Though it is widely appreciated that numerous TFs act in protein complexes to regulate cellular identity and function, the significance of their heterogenous expression for maintaining identity and function has not been explored. Building on the strength of scRNA-seq to resolve cell heterogeneity, we explored numerous islet-enriched TFs and found bimodal distribution patterns that suggest the presence of unique combinatorial profiles. In this manuscript, we investigated expression patterns of three TFs with known changes in islet cell development and diabetes: α cell-specific *ARX*, β cell-specific *MAFA*, and *MAFB*, which is expressed in both α and β cells and has a unique expression profile compared to rodent islets. Interestingly, other islet-enriched TFs were consistently elevated in *ARX*/*MAFB* co-expressing α cells and *MAFA*/*MAFB* co-expressing β cells, supporting the concept of islet-enriched TFs acting in self-regulating networks, and making it likely that combinatorial profiles of other TFs also reveal interesting populations with functional consequences. Larger datasets and network-based approaches considering additional TF combinations should be used to examine more complex expression patterns and how these patterns change in T1D and T2D islet cells.

Our data suggest that *ARX*/*MAFB* co-expressing α cells and *MAFA*/*MAFB* co-expressing β cells have elevated expression of functional genes compared to cells that express only one or neither factor. Nonetheless, elevated expression for certain genes in single TF-expressing populations (*e.g.*, *MDH2* and *KCNMA1* in *MAFB*-expressing β cells) may provide insight on how these individual TFs act in each cell type. Indeed, a comparison of our data to molecular studies of these TFs in mice or human stem cells reveals numerous similarities. For example, our data demonstrates that *MAFA*/*MAFB* co-expressing β cells are distinct from populations that express only a single TF which suggests that although these factors are related, they have distinct targets and roles within the β cell. This is consistent with a recent report showing that in mice, MAFB does not compensate for MAFA loss^12^. Further, our data highlights *MAFB* as playing a key role in defining both β and α cell identity, in line with a recent report where MAFB deletion in human embryonic stem cells disrupted the differentiation process for both β and α cells^55^. Thus, our approach highlights how transcription factor profiles at the single cell level can be used to predict transcriptional and functional consequences of genetic manipulation, highlighting an immense power for large scRNA-seq datasets.

While there were not sufficient cells for robust statistical comparison of all subsets, it is interesting to note that the electrophysiological profile of the cells expressing neither MAFA nor MAFB was similar to those cells expressing only one of the factors, thus suggesting a specific benefit to having combined expression of both factors in adult human β cells that is not apparent with only one of the TFs. These findings have several implications given the unique timing of MAFA and MAFB expression in the human β cell and differ slightly from our transcriptional data that suggested more of a progressive increase with double negative group showing the lowest expression, followed by single TF groups, and co-expressing cells having highest expression of genes related to hormone secretory function. Future investigation with larger functional datasets will be needed to further delineate these interesting findings as well as directly evaluate the role of MAFA, MAFB, and other enriched transcription factors in human islet cell hormone secretion.

One contribution to bimodal distribution of low-abundance transcripts like TFs is gene dropout, where a gene is detected only in a subset of cells due to low mRNA quantity. However, greater expression of functional genes in one subpopulation (often dual positive cells) suggests that dropout is not simply a stochastic event and could instead reflect cell activity or a biological process such as transcriptional bursting^56^. Further, we analyzed three additional scRNA-seq datasets of human islets generated by various single cell technologies^18, 28, 29^, and all showed trends consistent with the current study. Finally, taking advantage of the unique Patch-seq approach from our previous study, we were able to validate increased cellular function reflected by electrophysiological parameters (**Figure 7**). Together, these data indicate that our observations are not technical in nature and instead represent important aspects of human islet biology.

Given the potential inconsistencies between transcript and protein-level expression in human islets^53^, we pursued identification of heterogeneous TF protein expression in human pancreatic tissue. Though there were discrepancies in subpopulation distribution estimated by transcript versus by immunodetection, the presence of all TF combinations in tissue suggests this heterogeneity is not limited to one experimental approach. Differences may also arise from post-transcriptional control of protein levels that would not be apparent at the transcript level. Novel, single cell multi-omic techniques will be required to define the precise correlation between TF mRNA and protein abundance, and these techniques may also help define how the described heterogeneity relates to other forms of β cell heterogeneity that have been previously described or hypothesized^18–22^. Heterogeneity within α cell populations has been less studied but our data indicate it may have an unappreciated role within the islet as well.

There are limitations to the current study that suggest opportunities for future work. First, the dispersion of islet cells required for scRNA-seq disrupts the microenvironment, which is known to be crucial for coordinated islet function^57, 58^. How the α and β cell subpopulations defined in this study, function in the islet context is presently unknown – while having all highly functional cells would seem beneficial, some data has suggested that both mature and immature cells are required within an islet for optimal function^59^. Importantly, the nature of scRNA-seq means we cannot discern whether the heterogeneity described here is stable or a snapshot of a dynamic cell state.

In sum, we highlight the utility of a large, scRNA-seq dataset by uncovering previously unappreciated heterogeneity in combined key islet-enriched TF expression and demonstrate that it has implications for β cell function. Ultimately, defining the key characteristics of highly functional α and β cells will allow not only a greater understanding of pathways governing coordinated hormone secretion but also engineering of cells closely resembling native α or β cell function for cell replacement therapy to treat diabetes.

## MATERIALS AND METHODS

### Human pancreatic islet samples

Human islet preparations (n=5; see **Table S1** for donor information) were obtained through partnerships with the Integrated Islet Distribution Program (IIDP, RRID:SCR_014387; http://iidp.coh.org/), Alberta Diabetes Institute (ADI) IsletCore (RRID:SCR_018566; https://www.epicore.ualberta.ca/IsletCore/), and the Human Pancreas Analysis Program (HPAP, RRID:SCR_016202; https://hpap.pmacs.upenn.edu/) of the Human Islet Research Network.

Assessment of human islet function was performed by islet macroperifusion assay on the day of arrival, as previously described ^60^. Islets were cultured in CMRL 1066 media (5.5 mM glucose, 10% FBS, 1% Penicillin/Streptomycin, 2 mM L-glutamine) in 5% CO2 at 37°C for <24 hours prior to beginning studies. The Vanderbilt University Institutional Review Board does not consider deidentified human pancreatic specimens to qualify as human subject research.

This study used data from the Organ Procurement and Transplantation Network (OPTN) that was in part compiled from the Data Hub accessible to IIDP-affiliated investigators through IIDP portal (https://iidp.coh.org/secure/isletavail). The OPTN data system includes data on all donors, wait-listed candidates, and transplant recipients in the US, submitted by the members of the OPTN. The Health Resources and Services Administration (HRSA), U.S. Department of Health and Human Services provides oversight to the activities of the OPTN contractor. The data reported here have been supplied by UNOS as the contractor for the Organ Procurement and Transplantation Network (OPTN). The interpretation and reporting of these data are the responsibility of the author(s) and in no way should be seen as an official policy of or interpretation by the OPTN or the U.S. Government.

### Cell preparation

Handpicked pancreatic islets were dispersed by manual pipetting using 0.025% HyClone trypsin (Cytiva/GE Healthcare #SH30042.01) and subsequently quenched with RPMI media containing 20% FBS (Millipore #TMS-013-B). Cells were washed in the same media twice followed by one wash with 0.04% BSA (Thermo Scientific # AM2616) in 1X PBS without calcium and magnesium (Corning Cellgro #21-040-CV). Washed cells were immediately counted in a Trypan blue stain-based Cell Countess II Automated Cell Counter (Thermo Scientific #AMQAX1000).

Viability obtained from the cell preparations ranged from 70-85%. Cells were resuspended in 0.04% BSA/1X PBS at a density of 630-1,200 cells/μl in preparation for single cell RNA sequencing.

### Purification of α and β cell by FACS

Human islets from preparations #1 and 2 (**Table S1**) were dispersed and sorted for α and β cells following protocol described previously^17, 49, 61^. Briefly, 0.025% trypsin was used to disperse islet cells by manual pipetting and subsequently quenched with RPMI containing 10% FBS. Previously characterized primary and secondary antibodies^25, 27, 62^ are listed in **Table S3** and gating strategy is shown in **Figure S2A**. Collected α and β cells for scRNA-seq were washed in 1X PBS with 0.04% BSA and immediately loaded into the 10x Chromium Controller at 1,200 cells/μl based on FACS counts, with single cell libraries prepared as described below. In parallel, 10,000 α and β cells from islet preparation #2 were stored in RNA extraction buffer to be processed for bulk RNA-seq as described below.

### Bulk RNA library preparation and sequencing

RNA was extracted from sorted α and β cells using the Invitrogen RNAqueous-Micro Total RNA Isolation kit (Thermo Fisher #AM1931). TURBO DNA-free (Ambion) was used to treat any trace DNA contamination. RNA was quantified by Qubit Fluorometer 2.0 and RNA integrity was confirmed (RIN >7) by 2100 Bioanalyzer (Agilent). RNA was amplified using NuGen Ovation RNA amplification kit and sheared to an average size of 200 bp, then libraries were prepared using the NEBNext DNA library prep kit (New England Biolabs). Final libraries were sequenced on a Novaseq platform (Illumina), using paired-end reads (50 bp) targeting 50 million reads per sample. Raw reads were aligned to human reference genome hg38 using STAR v2.6^63^. Strand NGS 3.4 commercial software was used to import aligned files (.bam) and subsequently check alignment quality, filter reads based on read quality, quantify transcripts, and normalize counts to transcript per million (TPM). Only genes with expression log2 (TPM) > 1 were considered for the analysis in **Figures 2B-H** and **S2B-E**. Gene Ontology (GO) analyses were performed using enrichDAVID function of R package ClusterProfiler^64^ 3.14.3 (**Figures 2D-G**, **S2B-C**) or DAVID v6.8 web service^65^ (available https://david.ncifcrf.gov) (**Figure S2D-E**). Differential expression analysis between α and β cells was defined as fold change ≥±1, calculated based on p-value estimated by z-score calculations (cutoff 0.05) as determined by Benjamini Hochberg false discovery rate (FDR) correction of 0.05^66^.

For original/source data used in **Figures 1A and 1C,** bulk RNA-seq of sorted human islet α is available under NCBI GEO accession numbers GSE106148 (Brissova et al. 2018)^17^, bulk RNA-seq of sorted human islet β cells is available under GSE116559 (Saunders et al. 2019)^27^. For original/source data used in **Figure 1B, 1D, S1A, S1B**, bulk RNA-seq data is available under GSE57973 (Arda *et al*. 2016)^10^, and GSE67543 (Blodgett et al. 2015)^26^ utilizing publicly available Reads Per Kilobase Per Million (RPKM) and Transcripts Per Million (TPM) normalized counts from NCBI GEO respectively.

### Single cell library preparation and sequencing

Sorted or dispersed islet cell samples were loaded in triplicate (approximately 10,000 cells/replicate) on 10x Chromium chips (PN# 1000009) to ensure consistent results. Gel Bead in Emulsion (GEM) generation and barcoding were performed on the 10x Chromium Controller according to the manufacturer’s instructions (10x Genomics Single Cell 3’ Library and Gel bead Kit v2 #220104). Immediately after GEMs were generated, samples were transferred to a 0.2ml TempAssure PCR 8-tube strip (USA Scientific #14024700), capped, and placed into a thermocycler (Bio-Rad T100™ Thermal Cycler) for reverse transcription. After incubation, the GEMs were broken, and pooled cDNA proceeded to cleanup using Silane magnetic beads (10x Genomics #2000048) to remove leftover reagents. cDNA was then amplified through 10 cycles of PCR and cleaned using SPRIselect beads (Beckman Coulter # B23318). Resulting cDNA (average 45 ng/replicate) was checked for quality by Qubit dsDNA HS Assay Kit (Thermo Fisher Scientific #Q32854) and Agilent Bioanalyzer High Sensitivity Kit (Agilent #5067-4626). Final libraries were constructed according to manufacturer’s instruction and underwent 14 cycles of PCR amplification after sample index addition, yielding ∼953ng and average library size of 486bp. Final libraries were sequenced with a Novaseq sequencer (Illumina) using paired-end reads (100 bp) to average depth of ∼146,000 reads per cell.

### scRNA-seq alignment, preprocessing, and quality control

Alignment to reference transcriptome (GRCh38-1.2; gene annotation provided by 10x Genomics) and unique molecular identifier (UMI)-based gene expression quantification was obtained following the Cell Ranger analysis pipeline (v2.1). The “Aggr” function was used to aggregate transcript counts and normalize read depth across 5 islet preparations and their technical replicates, producing one single gene-cell (feature-barcode) matrix. In **Figure 3**, the “Aggr” function was applied to 2 islet preparations, including the samples that were FACS-sorted. Further data preprocessing and clustering was performed using Seurat version 3.1^50^.

Cells with 200-4,000 detected genes and <10% mitochondrial gene expression were retained, and only genes expressed in ≥3 cells were considered for further analysis. Gene expression was normalized for each cell by library size and log-transformed using a size factor of 10,000 molecules per cell. For feature selection, 2,000 highly variable genes were selected using function “FindVariableFeatures.” The data was further centered and scaled to zero mean and unit variance implemented in the “ScaleData” function using parameter “vars.to.regress” to regress out mitochondrial gene expression. Cells co-expressing the insulin (*INS*) and glucagon (*GCG*) genes above log expression of 6.5 and 5 respectively, as well as cells expressing *INS* or *GCG* in addition to any other cell type gene marker, were removed as doublets (see **Table S2** for cell type markers used). Transcript counts from lysed cells (ambient mRNA/background RNA) were estimated and genes identified from empty droplets (droplets without cells) using DropletUtils package^67^. Using the raw gene-barcode matrix (Cell Ranger v3.1), UMI threshold of 100 and below were considered ambient transcripts. About ∼200 genes were identified as ambient genes and their expression level was noted to remove from the original gene barcode matrix in order to account for transcript stemming from lysed cells. The principal component analysis (PCA) was performed using previously determined 2,000 high variable genes as input. An elbow plot, which ranks the principal components (PCs) based on percent variance per PC, was considered to determine the number of PCs to use for downstream graph-based clustering. “FindNeighbors” and “FindClusters” functions were used with 20 PCs as input for cluster generation and resolution at 0.6. Finally, UMAP dimension reduction was used for cluster visualization.

### Immunohistochemical analysis

Lightly paraformaldehyde (PFA)-fixed human pancreatic tissue cryosections from n=3 donors (age range 20-55 years) were prepared for immunohistochemistry and stained as described previously^17, 27, 61^. Primary and secondary antibodies and their dilutions are listed in **Table S3**; donor information is supplied in **Table S4**. Images were acquired at 20X with 2X digital zoom using a FV3000 confocal laser scanning microscope (Olympus) and processed using HALO software (Indica Labs) with a cytonuclear algorithm (HighPlex FL v3.2.1) to set an intensity threshold (“hi/lo”) for each marker.

### Analysis of previously published scRNA-seq datasets

Raw gene count matrices were extracted from existing single cell RNA-seq datasets^18, 28, 29^ and further analyzed using the R package Seurat version 3.1 as described above.

### Single cell electrophysiology and gene expression

Patch-seq was performed as described previously in Camunas et al.^18^.

### Statistical Information

Specific statistical tests used for each dataset are described in the figure legends and text where appropriate. Pearson’s correlation (**Figures 2B-C** and **3C-D**) was performed using the ‘ggpubr’ package (available http://rpkgs.datanovia.com/ggpubr/). Gene clustering analysis (**Figure 2D-G**) was performed using the “enrichDAVID” function of the R package clusterProfiler 3.14.3^64^. All other statistical analyses were performed using GraphPad Prism software.

## DATA AVAILABILITY

Sequencing datasets GEO.

Single cell dataset visualization. Access from: https://powersbrissovalab.shinyapps.io/scRNAseq-Islets/

## AUTHOR CONTRIBUTIONS

Conceptualization, S.S., D.C.S., J.T.W., M.B., R.S., and A.C.P.; Methodology, S.S., D.C.S., J.T.W., J. C.-S., and X.-Q.D.; Investigation, S.S., D.C.S., J.T.W., X.-Q.D., R.H., R.A., G.P., and R.B.; Formal Analysis, S.S., J.C.-S., and J.-P.C.; Writing – Original Draft, S.S., D.C.S., and J.T.W.; Writing – Review & Editing, all authors; Funding Acquisition, P.E.M., S.E.L., A.C.P., and M.B.; Supervision, N.P., J.-P.C., S.C.J.P., P.E.M., S.E.L., A.C.P., and M.B.

## ACKNOWLEDGMENT

We thank the organ donors and their families for their invaluable donations and the International Institute for Advancement of Medicine (IIAM), Organ Procurement Organizations, National Disease Research Exchange (NDRI), and the Alberta Diabetes Institute IsletCore together with the Human Organ Procurement and Exchange (HOPE) program and Trillium Gift of Life Network (TGLN) for their partnership in studies of human pancreatic tissue for research. This study used human pancreatic islets that were provided by the NIDDK-funded Integrated Islet Distribution Program at the City of Hope (NIH Grant # 2UC4 DK098085). This work was supported by the Human Islet Research Network (HIRN; RRID:SCR_014393; https://hirnetwork.org; DK112232, DK123716, DK123743, DK120447, DK120456, DK104211, DK108120, DK090570), and by DK106755, DK117147, T32GM007347, F30DK118830, DK20593 (Vanderbilt Diabetes Research and Training Center), The Leona M. and Harry B. Helmsley Charitable Trust, the JDRF, and the Department of Veterans Affairs (BX000666). Flow cytometry was performed in the Vanderbilt Flow Cytometry Shared Resource (P30 CA68485, DK058404).

## REFERENCES

1. Noguchi, G. M. & Huising, M. O. Integrating the inputs that shape pancreatic islet hormone release. Nat Metabolism 1, 1189–1201 (2019).

2. Chen, C., Cohrs, C. M., Stertmann, J., Bozsak, R. & Speier, S. Human beta cell mass and function in diabetes: Recent advances in knowledge and technologies to understand disease pathogenesis. Mol Metab 6, 943–957 (2017).

3. Cnop, M. et al. Mechanisms of Pancreatic β-Cell Death in Type 1 and Type 2 Diabetes: Many Differences, Few Similarities. Diabetes 54, S97–S107 (2005).

4. Halban, P. A. et al. β-Cell Failure in Type 2 Diabetes: Postulated Mechanisms and Prospects for Prevention and Treatment. Diabetes Care 37, 1751–1758 (2014).

5. Unger, R. H. & Cherrington, A. D. Glucagonocentric restructuring of diabetes: a pathophysiologic and therapeutic makeover. J Clin Invest 122, 4–12 (2012).

6. Pan, F. C. & Wright, C. Pancreas organogenesis: From bud to plexus to gland. Dev Dynam 240, 530 565 (2011).

7. Jennings, R. E., Berry, A. A., Strutt, J. P., Gerrard, D. T. & Hanley, N. A. Human pancreas development. Development 142, 3126–3137 (2015).

8. Zhu, Z. et al. Genome Editing of Lineage Determinants in Human Pluripotent Stem Cells Reveals Mechanisms of Pancreatic Development and Diabetes. Cell Stem Cell 18, 755–68 (2016).

9. Thompson, P. & Bhushan, A. β Cells led astray by transcription factors and the company they keep. J Clin Invest 127, 94 97 (2017).

10. Arda, H. E. et al. Age-Dependent Pancreatic Gene Regulation Reveals Mechanisms Governing Human β Cell Function. Cell Metab 23, 909–920 (2016).

11. Dai, C. et al. Islet-enriched gene expression and glucose-induced insulin secretion in human and mouse islets. Diabetologia 55, 707–718 (2012).

12. Cyphert, H. A. et al. Examining How the MAFB Transcription Factor Affects Islet β Cell Function Postnatally. Diabetes 68, db180903 (2018).

13. Hang, Y. et al. The MafA Transcription Factor Becomes Essential to Islet β-Cells Soon After Birth. Diabetes 63, 1994–2005 (2014).

14. Guo, S. et al. Inactivation of specific β cell transcription factors in type 2 diabetes. J Clin Invest 123, 3305–3316 (2013).

15. Dai, C. et al. Stress-impaired transcription factor expression and insulin secretion in transplanted human islets. J Clin Invest 126, 1857–1870 (2016).

16. Talchai, C., Xuan, S., Lin, H. V., Sussel, L. & Accili, D. Pancreatic β cell dedifferentiation as a mechanism of diabetic β cell failure. Cell 150, 1223–1234 (2012).

17. Brissova, M. et al. α Cell Function and Gene Expression Are Compromised in Type 1 Diabetes. Cell Reports 22, 2667–2676 (2018).

18. Camunas-Soler, J. et al. Patch-Seq Links Single-Cell Transcriptomes to Human Islet Dysfunction in Diabetes. Cell Metab 31, 1017–1031.e4 (2020).

19. Dorrell, C. et al. Human islets contain four distinct subtypes of β cells. Nat Commun 7, 11756 (2016).

20. Wang, Y. J. et al. Single-Cell Mass Cytometry Analysis of the Human Endocrine Pancreas. Cell Metab 24, 616–626 (2016).

21. Thompson, P. J. et al. Targeted Elimination of Senescent Beta Cells Prevents Type 1 Diabetes. Cell Metab 29, 1045–1060.e10 (2019).

22. Meulen, T. van der & Huising, M. O. Maturation of Stem Cell-Derived Beta-cells Guided by the Expression of Urocortin 3. Rev Diabet Stud 11, 115–132 (2014).

23. Fadista, J. et al. Global genomic and transcriptomic analysis of human pancreatic islets reveals novel genes influencing glucose metabolism. Proc National Acad Sci 111, 13924–13929 (2014).

24. Eizirik, D. L. et al. The Human Pancreatic Islet Transcriptome: Expression of Candidate Genes for Type 1 Diabetes and the Impact of Pro-Inflammatory Cytokines. Plos Genet 8, e1002552 (2012).

25. Dorrell, C. et al. Transcriptomes of the major human pancreatic cell types. Diabetologia 54, 2832–2844 (2011).

26. Blodgett, D. M. et al. Novel Observations From Next-Generation RNA Sequencing of Highly Purified Human Adult and Fetal Islet Cell Subsets. Diabetes 64, 3172–3181 (2015).

27. Saunders, D. C. et al. Ectonucleoside Triphosphate Diphosphohydrolase-3 Antibody Targets Adult Human Pancreatic β Cells for In Vitro and In Vivo Analysis. Cell Metab 29, 745–754.e4 (2019).

28. Baron, M. et al. A Single-Cell Transcriptomic Map of the Human and Mouse Pancreas Reveals Inter- and Intra-cell Population Structure. Cell Syst 3, 346–360.e4 (2016).

29. Segerstolpe, Å. et al. Single-Cell Transcriptome Profiling of Human Pancreatic Islets in Health and Type 2 Diabetes. Cell Metab 24, 593–607 (2016).

30. Xin, Y. et al. RNA Sequencing of Single Human Islet Cells Reveals Type 2 Diabetes Genes. Cell Metab 24, 608–615 (2016).

31. Lawlor, N. et al. Single-cell transcriptomes identify human islet cell signatures and reveal cell-type-specific expression changes in type 2 diabetes. Genome Res 27, 208–222 (2017).

32. Fang, Z. et al. Single-Cell Heterogeneity Analysis and CRISPR Screen Identify Key β-Cell-Specific Disease Genes. Cell Reports 26, 3132–3144.e7 (2019).

33. Wang, Y. J. & Kaestner, K. H. Single-Cell RNA-Seq of the Pancreatic Islets--a Promise Not yet Fulfilled? Cell Metab 29, 539–544 (2019).

34. Mawla, A. M. & Huising, M. O. Navigating the Depths and Avoiding the Shallows of Pancreatic Islet Cell Transcriptomes. Diabetes 68, 1380–1393 (2019).

35. Itoh, M. et al. Partial loss of pancreas endocrine and exocrine cells of human ARX-null mutation: Consideration of pancreas differentiation. Differentiation 80, 118–122 (2010).

36. Gosmain, Y., Cheyssac, C., Masson, M. H., Dibner, C. & Philippe, J. Glucagon gene expression in the endocrine pancreas: the role of the transcription factor Pax6 in α-cell differentiation, glucagon biosynthesis and secretion. Diabetes Obes Metabolism 13, 31–38 (2011).

37. Courtney, M. et al. The Inactivation of Arx in Pancreatic α-Cells Triggers Their Neogenesis and Conversion into Functional β-Like Cells. Plos Genet 9, e1003934 (2013).

38. Wang, H., Brun, T., Kataoka, K., Sharma, A. J. & Wollheim, C. B. MAFA controls genes implicated in insulin biosynthesis and secretion. Diabetologia 50, 348 358 (2007).

39. Bonnavion, R. et al. Both PAX4 and MAFA Are Expressed in a Substantial Proportion of Normal Human Pancreatic Alpha Cells and Deregulated in Patients with Type 2 Diabetes. Plos One 8, e72194 (2013).

40. Liu, W. et al. Abnormal regulation of glucagon secretion by human islet alpha cells in the absence of beta cells. Ebiomedicine (2019) doi:10.1016/j.ebiom.2019.11.018.

41. Matsuoka, T. et al. The MafA transcription factor appears to be responsible for tissue-specific expression of insulin. P Natl Acad Sci Usa 101, 2930–2933 (2004).

42. Matsuoka, T. et al. MafA Regulates Expression of Genes Important to Islet β-Cell Function. Mol Endocrinol 21, 2764–2774 (2007).

43. Artner, I. et al. MafA and MafB regulate genes critical to beta-cells in a unique temporal manner. Diabetes 59, 2530–2539 (2010).

44. Otonkoski, T., Andersson, S., Knip, M. & Simell, O. Maturation of Insulin Response to Glucose During Human Fetal and Neonatal Development: Studies with Perifusion of Pancreatic Isletlike Cell Clusters. Diabetes 37, 286–291 (1988).

45. Henquin, J.-C. & Nenquin, M. Dynamics and Regulation of Insulin Secretion in Pancreatic Islets from Normal Young Children. Plos One 11, e0165961 (2016).

46. Helman, A. et al. A Nutrient-Sensing Transition at Birth Triggers Glucose-Responsive Insulin Secretion. Cell Metab 31, 1004–1016.e5 (2020).

47. Matsuoka, T. et al. Members of the Large Maf Transcription Family Regulate Insulin Gene Transcription in Islet β Cells. Mol Cell Biol 23, 6049–6062 (2003).

48. Dorrell, C. et al. Isolation of major pancreatic cell types and long-term culture-initiating cells using novel human surface markers. Stem Cell Res 1, 183–194 (2008).

49. Saunders, D. C. et al. Ectonucleoside Triphosphate Diphosphohydrolase-3 Antibody Targets Adult Human Pancreatic β Cells for In Vitro and In Vivo Analysis. Cell Metab 29, 745–754.e4 (2019).

50. Butler, A., Hoffman, P., Smibert, P., Papalexi, E. & Satija, R. Integrating single-cell transcriptomic data across different conditions, technologies, and species. Nat Biotechnol 36, 411 (2018).

51. Stuart, T. et al. Comprehensive Integration of Single-Cell Data. Cell 177, 1888–1902.e21 (2019).

52. McInnes, L., Healy, J., Saul, N. & Großberger, L. UMAP: Uniform Manifold Approximation and Projection. J Open Source Softw 3, 861 (2018).

53. . Wortham, M. et al. Integrated In Vivo Quantitative Proteomics and Nutrient Tracing Reveals Age-Related Metabolic Rewiring of Pancreatic β Cell Function. Cell Reports 25, 2904–2918.e8 (2018).

54. Gustafsson, J. et al. Sources of variation in cell-type RNA-Seq profiles. Plos One 15, e0239495 (2020).

55. Russell, R. et al. Loss of the transcription factor MAFB limits β-cell derivation from human PSCs. Nat Commun 11, 2742 (2020).

56. Kharchenko, P. V., Silberstein, L. & Scadden, D. T. Bayesian approach to single-cell differential expression analysis. Nat Methods 11, 740–742 (2014).

57. Elliott, A. D., Ustione, A. & Piston, D. W. Somatostatin and insulin mediate glucose-inhibited glucagon secretion in the pancreatic α-cell by lowering cAMP. Am J Physiol-endoc M 308, E130–E143 (2015).

58. Capozzi, M. E. et al. β-Cell tone is defined by proglucagon peptides through cyclic AMP signaling. Jci Insight 4, e126742 (2019).

59. Nasteska, D. et al. PDX1LOW MAFALOW β-cells contribute to islet function and insulin release. Nat Commun 12, 674 (2021).

60. Kayton, N. S. et al. Human islet preparations distributed for research exhibit a variety of insulin-secretory profiles. Am J Physiol-endoc M 308, E592–E602 (2015).

61. Haliyur, R. et al. Human islets expressing HNF1A variant have defective β cell transcriptional regulatory networks. J Clin Invest 46, 1081–1087 (2018).

62. Bramswig, N. C. et al. Epigenomic plasticity enables human pancreatic α to β cell reprogramming. J Clin Invest 123, 1275–1284 (2013).

63. Dobin, A. et al. STAR: ultrafast universal RNA-seq aligner. Bioinformatics 29, 15–21 (2013).

64. Yu, G., Wang, L.-G., Han, Y. & He, Q.-Y. clusterProfiler: an R Package for Comparing Biological Themes Among Gene Clusters. Omics J Integr Biology 16, 284–287 (2012).

65. Huang, D. W. et al. Extracting biological meaning from large gene lists with DAVID. in undefined (eds. Baxevanis, A. D., Petsko, G. A., Stein, L. D. & Stormo, G. D.) vol. Chapter 13 Unit 11 (2009).

66. Benjamini, Y. & Hochberg, Y. Controlling the False Discovery Rate: A Practical and Powerful Approach to Multiple Testing. J Royal Statistical Soc Ser B Methodol 57, 289–300 (1995).

67. Lun, A. T. L. et al. EmptyDrops: distinguishing cells from empty droplets in droplet-based single-cell RNA sequencing data. Genome Biol 20, 63 (2019).

