## Supplemental Information for "Combinatorial transcription factor profiles predict mature and functional human islet α and β cells"

#### SUPPLEMENTAL FIGURE LEGENDS

**Supplemental Figure 1. ARX is expressed specifically in human α cells and MAFA in β cells, while MAFB is expressed in both α and β cells. (A–B)** Normalized expression of *ARX*, *MAFA*, and *MAFB* in previously published bulk RNA-sequencing (RNA-seq) datasets Arda *et al*. 2016[10] (**A**) and Blodgett *et al*. 2015[24] (**B**) from α cells (green) and β cells (blue). Bars in both panels show mean + SEM; symbols represent individual donors. Asterisks indicate significantly different (adjusted p-value <0.05) fold change (α vs. β). See also **Figure 1B**. **(C–D)** Immunohistochemical staining of pancreatic sections from nondiabetic adults (**Table S4**), showing specificity of ARX, MAFA, and MAFB (red) in α cells (GCG; green) and β cells (CPEP; blue). Arrowheads indicate cells negative (white) or positive (purple) for transcription factors; scale bars, 50 µm. See also **Figure 1C**.

**Supplemental Figure 2. Purified α and β cells analyzed by single cell or bulk RNA-sequencing. (A)** Gating strategy for sorted α and β cells identified by cell surface markers. Cell debris were excluded by forward scatter (FSC) and side scatter (SSC), single cells were identified by voltage pulse geometry (FSC-A v. FSC-H), and non-viable cells were excluded using propidium iodide (PI). Endocrine cell subpopulations were then isolated based on positivity for HPi1 (pan-endocrine marker) and additional positivity for HPa3 (α cells) or NTPDase3 (β cells). Antibody information can be found in **Table S3**. **(B–E)** Gene ontology enrichment of sorted α (**B**, **D**) and β (**C**, **E**) cells profiled by bulk and scRNA-seq (see schematic in **Figure 2A**). Analysis was run on genes detected by both scRNA-seq and bulk RNA-seq (“Common SC & Bulk;” n=1,076 for α cells and 797 for β cells), as well as the 2,000 most highly expressed genes in bulk samples not found in single cell samples (“Top 2,000 Unique Bulk”) using “enrichDAVID” function of clusterProfiler 3.14.3 R package. Colors correspond to populations depicted in **Figure 2,** B–C. In panel **C**, term with asterisk (*) is abbreviated due to space constraints; full term is ‘Positive regulation of transcription from RNA polymerase II promoter involved in cellular response to chemical stimulus.’

**Supplemental Figure 3. UMAP visualization of single cells identified with cell surface markers and those identified by unsupervised clustering. (A)** From each of two donors, α cells (WI-SC-α; pink) and β cells (WI-SC-β; green) were identified by unsupervised clustering following scRNA-seq of dispersed whole islets. **(B)** Cells positive for surface markers HPi1 and HPa3 (FACS-SC-α; teal) were enriched for α cells. **(C)** Cells positive for HPi1 and NTPDase3 (FACS-SC-β; purple) were enriched for β cells. Gating strategy for cells in **B**-**C** is shown in **Figure S2A**. **(D)** Number of genes detected was similar among α and β cells identified by cell surface markers and unsupervised clustering. See **Figure 2A** for schematic of experimental design.

**Supplemental Figure 4. Gene expression profiles of α and β cells identified by unsupervised clustering are highly concordant with those identified by cell surface markers. (A)** Principal component analysis (PCA) of cells identified by unsupervised clustering (WI-SC-α and WI-SC-β) overlaid with those obtained by fluorescence-activated cell sorting (FACS-SC-α and FACS-SC-β) in n=2 islet preparations (experimental schematic in **Figure 2A**). Data from both preparations combined in **Figure 3A**. **(B)** Heatmap of genes most highly contributing to variability in PCA; see also **Figure 3B**.

**Supplemental Figure 5. Donor-specific gene signatures of α and β cells identified by unsupervised clustering versus cell surface markers. (A–B)** Comparison of average log gene expression in α (**A**) and β (**B**) cells identified by unsupervised clustering (WI-SC-α and WI-SC-β) or cell surface markers (FACS-SC-α and FACS-SC-β) in n=2 islet preparations (experimental schematic in **Figure 2A**). Genes highlighted are α cell-enriched (yellow), β cell-enriched (blue), or selected markers of cell stress (grey). See also **Figure 3C-D**. **(B)** Heatmap showing variable expression of known α and β cell-enriched markers within and between each sample; see also **Figure 3E**.

**Supplemental Figure 6. Detailed characterization of endocrine cells from five nondiabetic human islet donors. (A)** Insulin and glucagon secretion were assessed in islets isolated from n=5 donors (age range 14–66 years) stimulated with 5.6 mM glucose (G 5.6), 16.7 mM glucose (G 16.7), 16.7 mM glucose + 100 mM isobutylmethylxanthine (IBMX) (G 16.7 + IBMX 100), 1.7 mM glucose + 1 mM epinephrine (G 1.7 + Epi 1), and 20 mM potassium chloride (KCl 20). Insulin and glucagon secretion is normalized to overall islet cell volume (expressed as islet equivalents; IEQs). **(B)** Bar graph illustrating cell type distribution within each islet preparation as per cell types annotated in **Figure 4A**. **(C)** UMAP of only α or β cells, showing clustering by islet preparation. See also **Figures 5A** and **6A**. **(D)** User-friendly web portal application for searching and viewing pancreatic cell types and their gene expression; available at https://shr19818.shinyapps.io/scRNAseq-Islets-v2/.

**Supplemental Figure 7. Raw expression values for transcription factor, α cell-enriched, ion channel, glucose metabolism, vesicle trafficking, exocytosis, and stress genes in *ARX*/*MAFB* populations.** Violin plots depict gene expression in α cell populations lacking *ARX* and *MAFB* (grey) and those expressing *MAFB* only (red), *ARX* only (blue), or co-expressing *both MAFB* and *ARX* (purple); n=24,248 total α cells. Data corresponds to dot plot in **Figure 5C**. Symbols underneath gene names indicate significance ($p<0.05$) from Tukey's multiple comparisons test following 2-way ANOVA.

**Supplemental Figure 8. Validation of α cell populations based on *ARX* and *MAFB* expression, as determined by previous scRNA-seq studies. (A)** Dot plots showing the expression patterns of selected genes related to cell identify, ion flux, glucose metabolism, vesicle trafficking, and exocytotic machinery. α cell populations (rows) are identified by expression of neither *ARX* nor *MAFB* (None), *ARX* only (ARX), *MAFB* only (MAFB), and co-expression of *ARX* and *MAFB* (Both). Dot size indicates the percentage of cells with detectable transcripts; color indicates gene's average scaled expression. Headers list study details from previously published datasets (Segerstolpe *et al.*, 2016[21]; Baron *et al.*, 2016[20]; Camunas-Soler *et al.*, 2020[15]) and final dot plot is as shown in **Figure 5C** for comparison. **(B)** Immunohistochemical staining of ARX (blue) and MAFB (red) in glucagon (GCG)-expressing α cells (green) of two nondiabetic adults (**Table S4**). Numbered arrowheads indicate the presence of α cell populations: 1, $ARX^{lo}$ $MAFB^{lo}$; 2, $ARX^{hi}$ $MAFB^{lo}$; 3, $ARX^{lo}$ $MAFB^{hi}$; 4, $ARX^{hi}$ $MAFB^{hi}$. See also **Figure 5E**.

**Supplemental Figure 9. Raw expression values for transcription factor, β-enriched, ion channel, glucose metabolism, vesicle trafficking, exocytosis, and stress genes in *MAFA*/*MAFB* populations.** Violin plots depict gene expression in β cell populations (n=11,034 total β cells) lacking *MAFA* and *MAFB* (grey) and those expressing *MAFA* only (red), *MAFB only* (blue), or co-expressing *both MAFA* and *MAFB* (purple); n=11,034 total β cells. Data corresponds to dot plot in **Figure 6C**. Symbols underneath gene names indicate significance ($p<0.05$) from Tukey's multiple comparisons test following 2-way ANOVA.

**Supplemental Figure 10. Validation of β cell populations based on *MAFA* and *MAFB* expression, as determined by previous scRNA-seq studies. (A)** Dot plots showing the expression patterns of selected genes related to cell identify, ion flux, glucose metabolism, vesicle trafficking, and exocytotic machinery. β cell populations (rows) are identified by expression of neither *MAFA* nor *MAFB* (None), *MAFA* only (MAFA), *MAFB* only (MAFB), and co-expression of *MAFA* and *MAFB* (Both). Dot size indicates the percentage of cells with detectable transcripts; color indicates gene's average scaled expression. Headers list study details from previously published datasets (Segerstolpe *et al.*, 2016[21]; Baron *et al.*, 2016[20]; Camunas-Soler *et al.*, 2020[15]) and final dot plot is as shown in **Figure 6C** for comparison. **(B)** Immunohistochemical staining of MAFA (red) and MAFB (blue) in C-peptide (CPEP)-expressing β cells (green) of two nondiabetic adults (**Table S4**). Numbered arrowheads indicate the presence of β cell populations: 1, MAFA$^{lo}$ MAFB$^{lo}$; 2, MAFA$^{hi}$ MAFB$^{lo}$; 3, MAFA$^{lo}$ MAFB$^{hi}$; 4, MAFA$^{hi}$ MAFB$^{hi}$. See also **Figure 6E**.

### Supplemental Figure 1

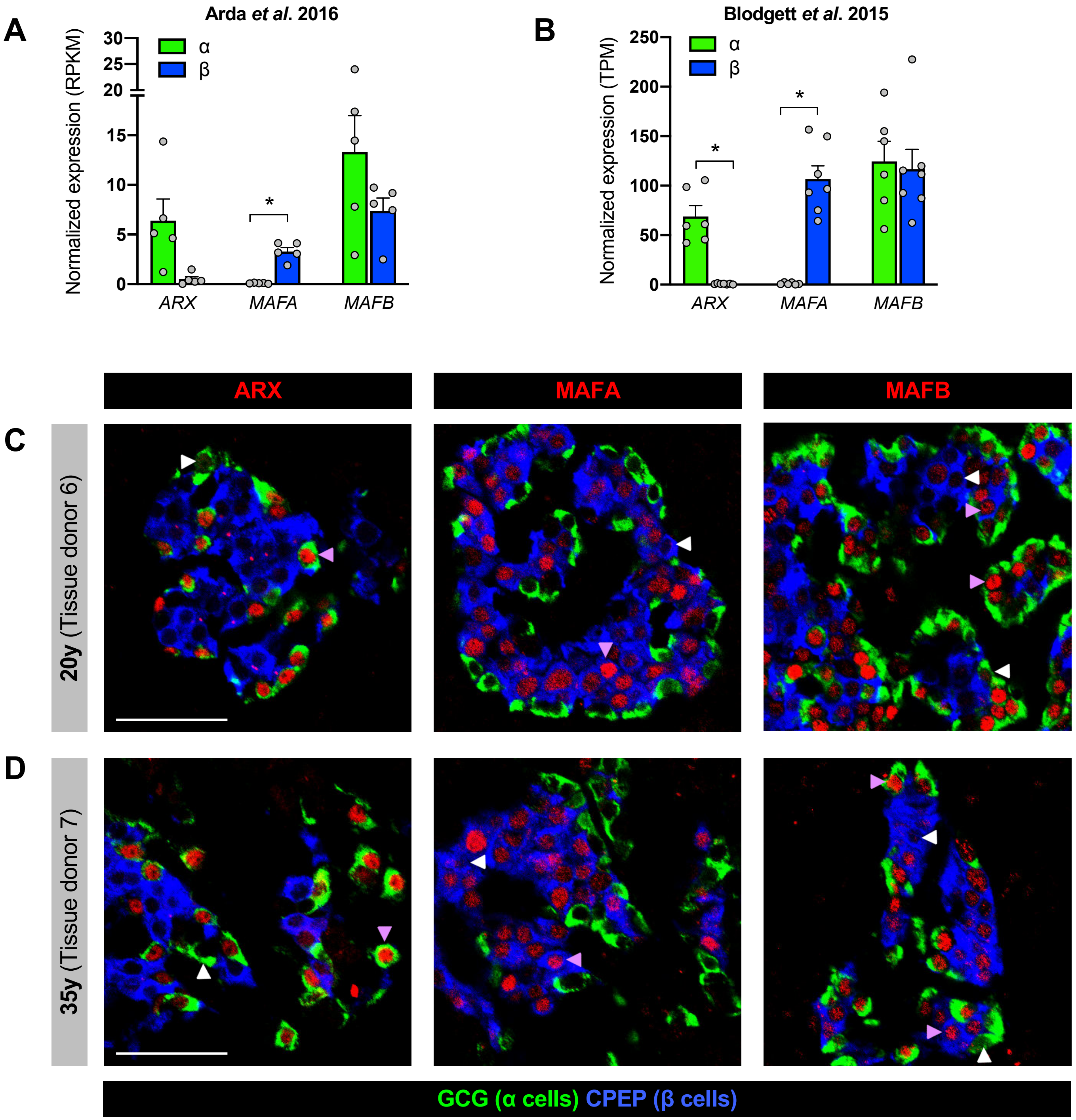

### Supplemental Figure 2

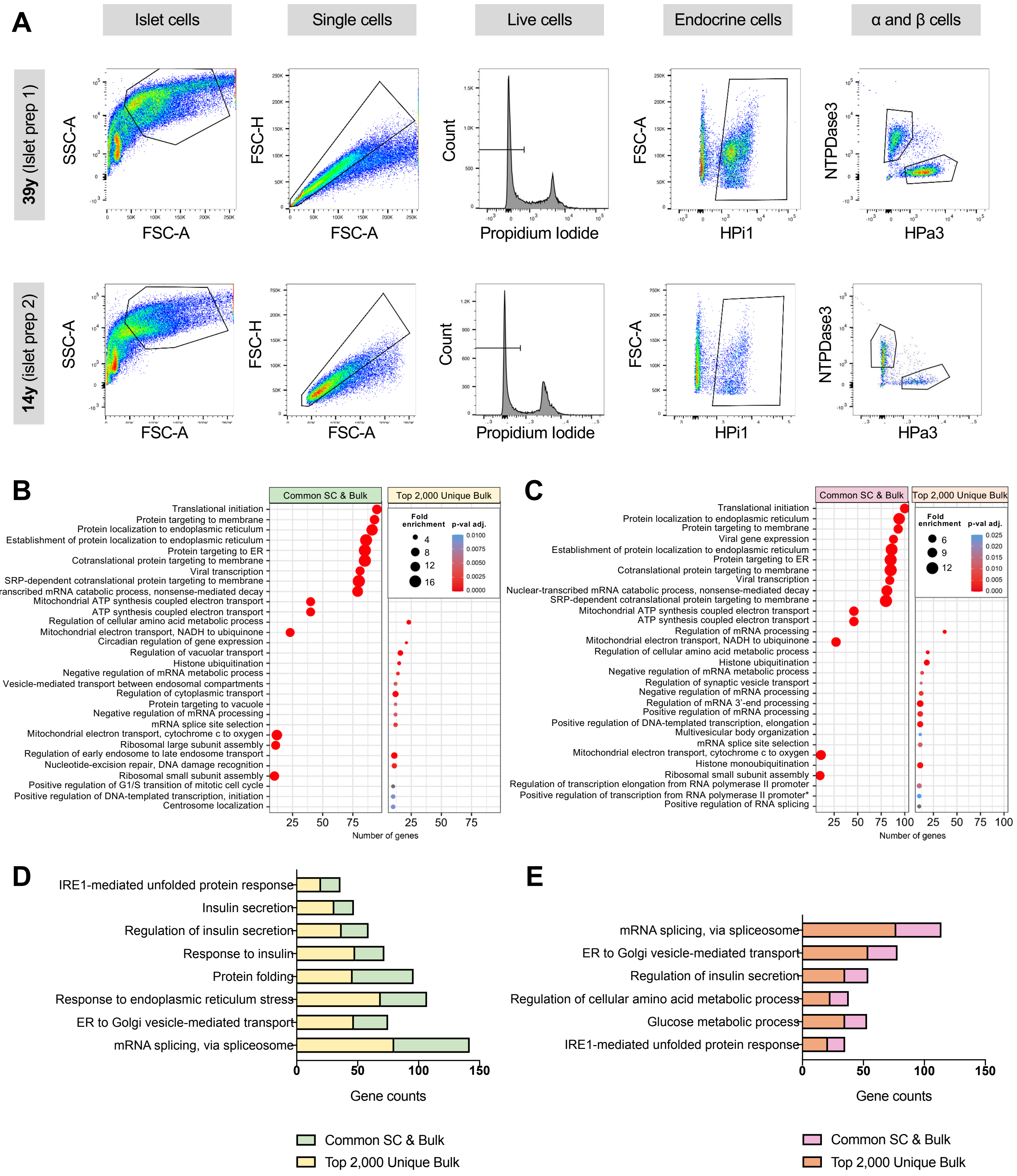

### Supplemental Figure 3

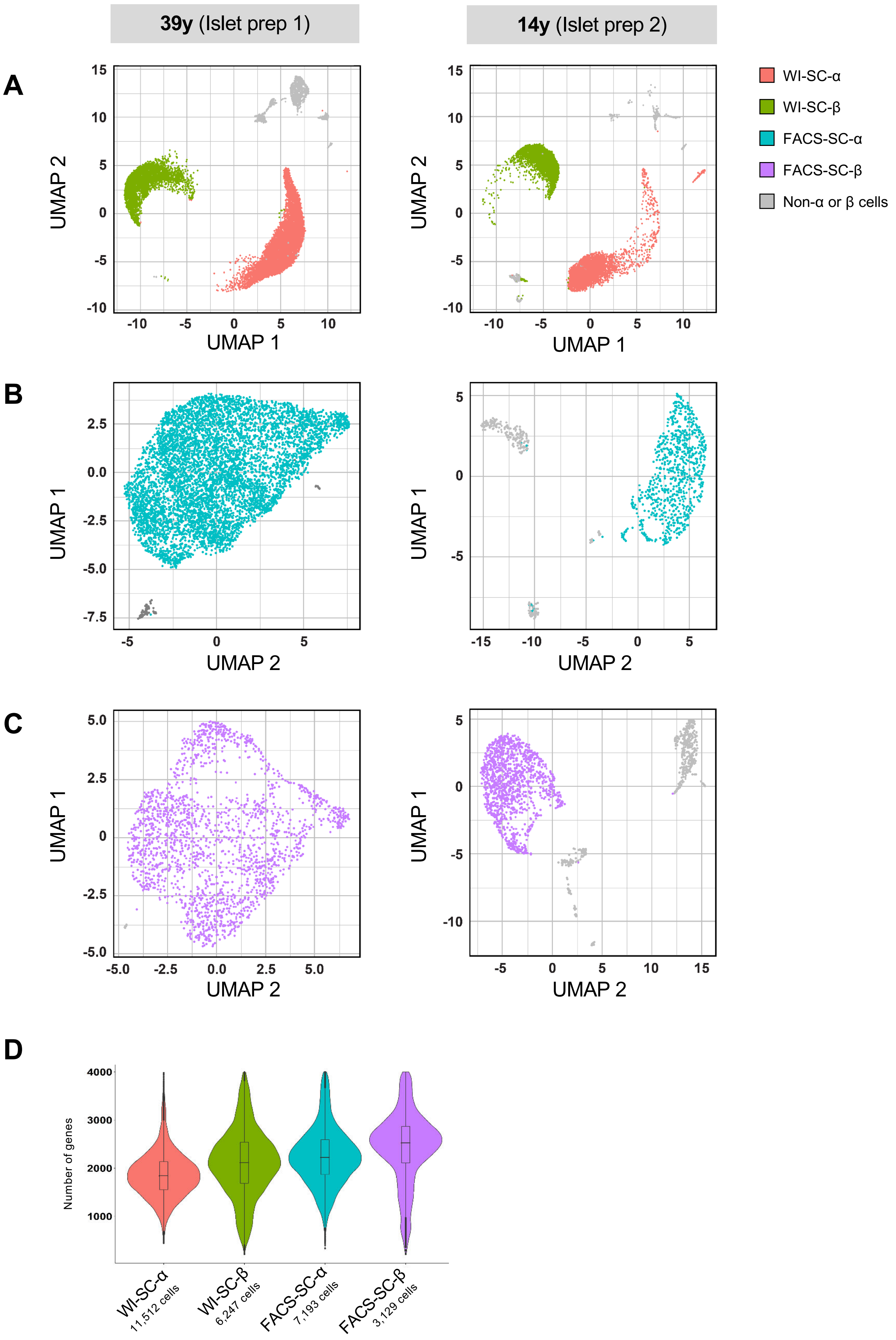

### Supplemental Figure 4

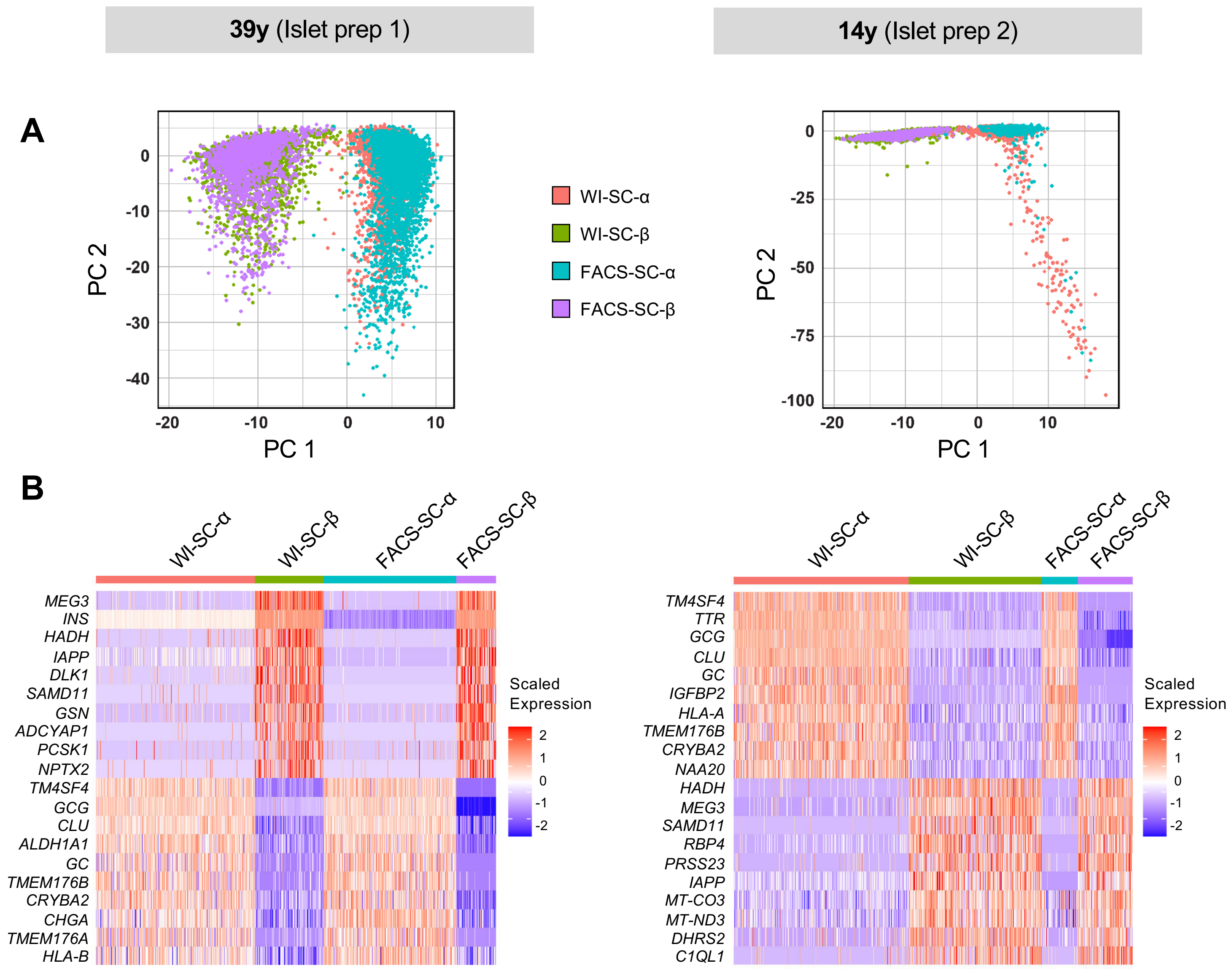

#### Supplemental Figure 5

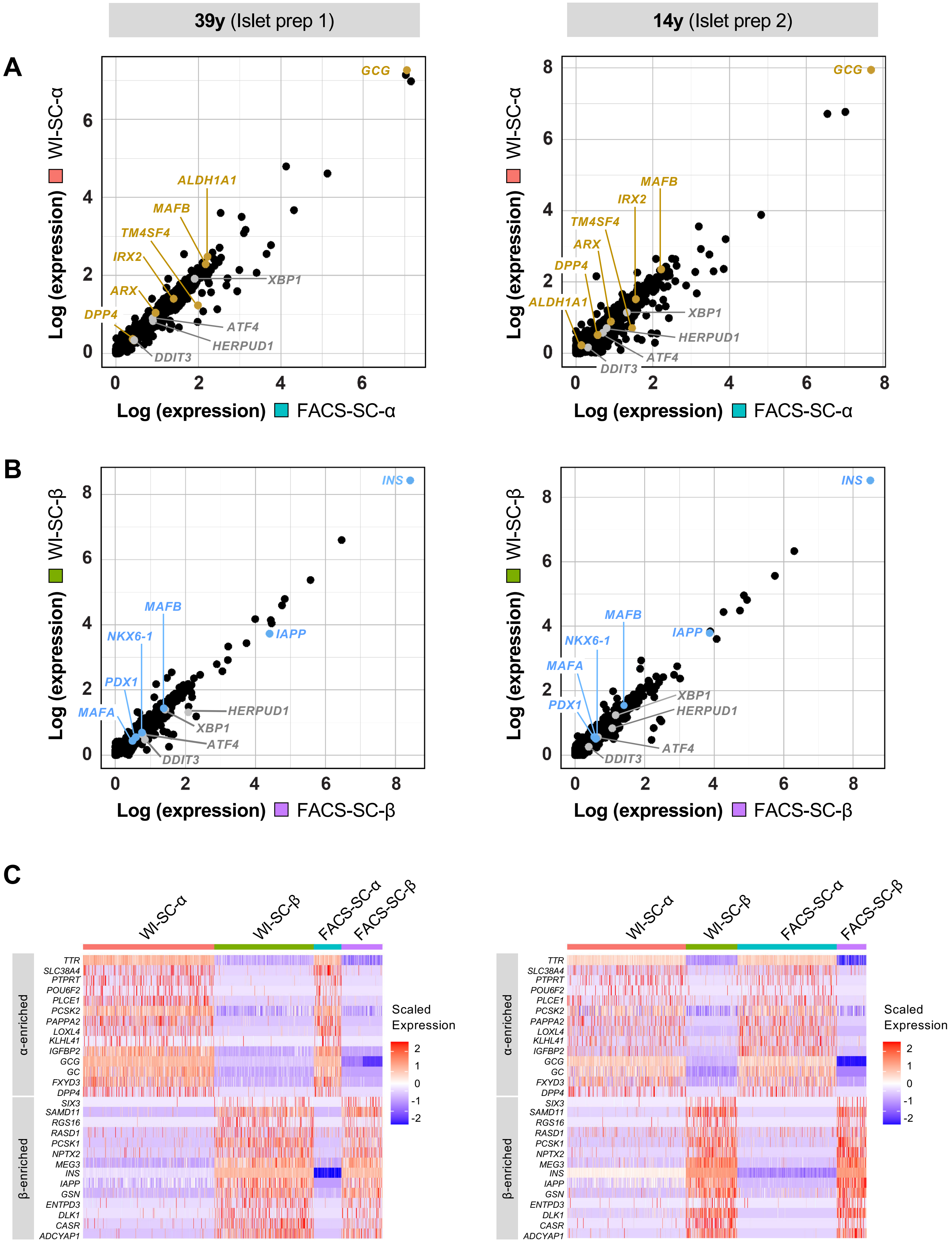

### Supplemental Figure 6

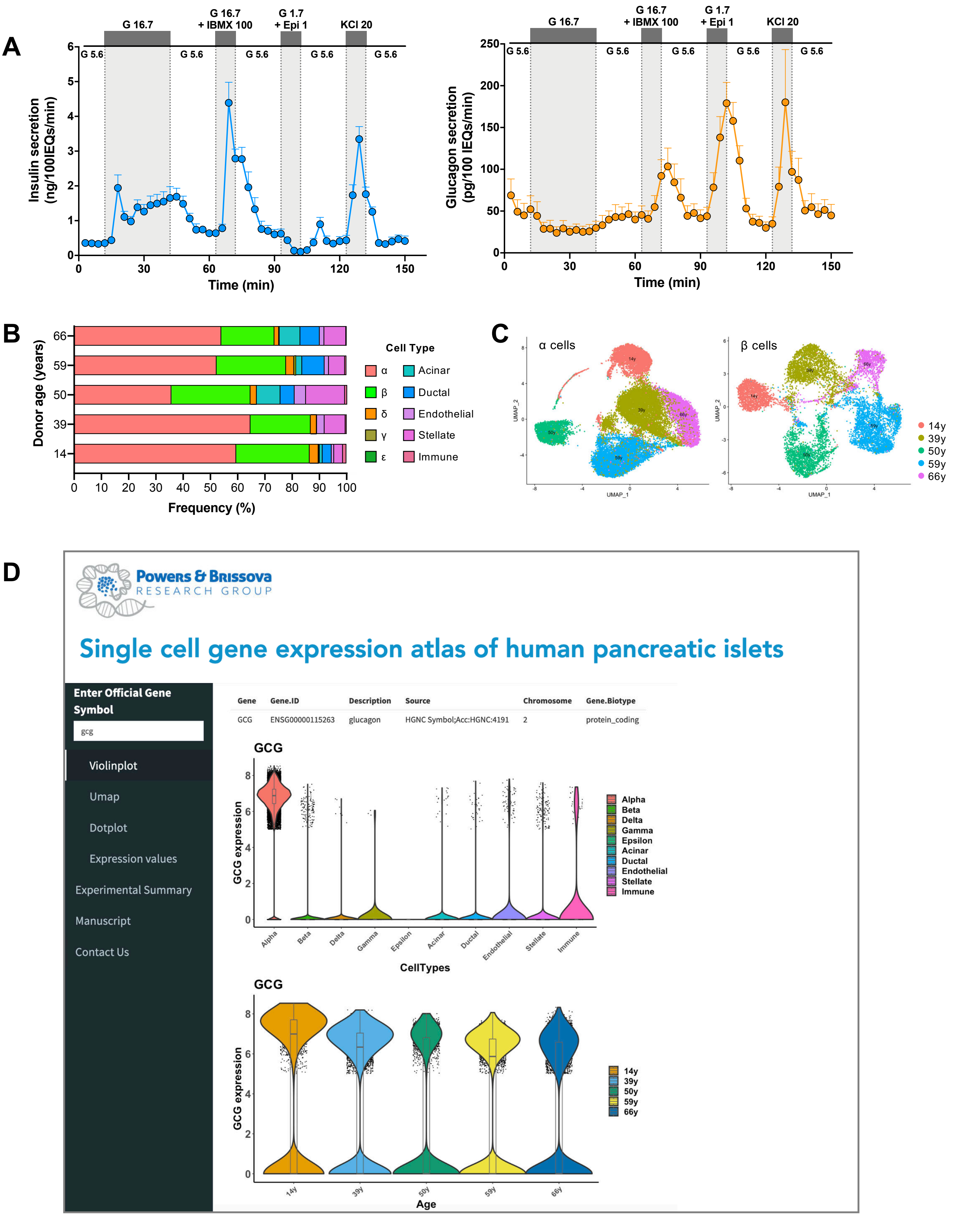

### Supplemental Figure 7

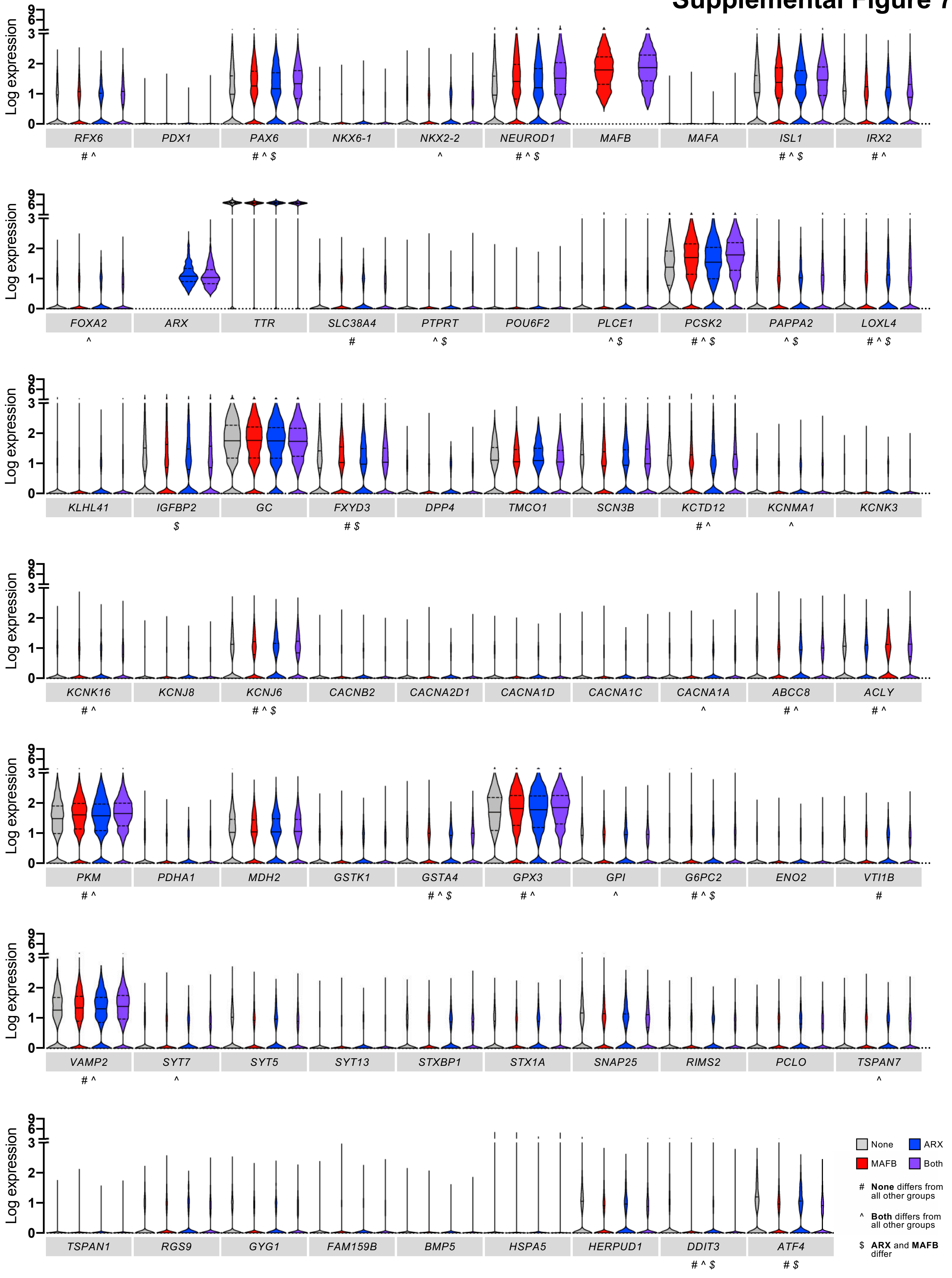

### Supplemental Figure 8

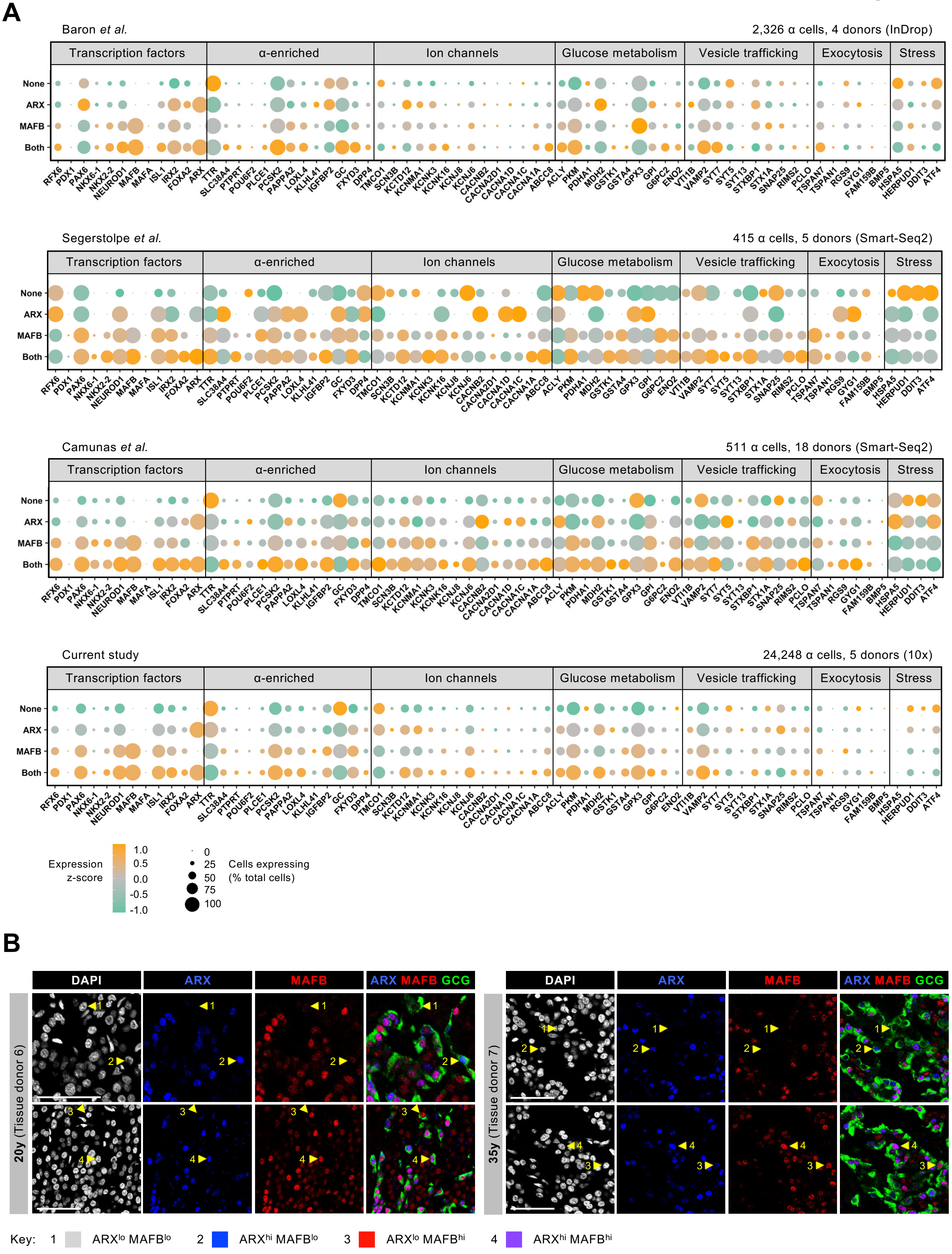

#### Supplemental Figure 9

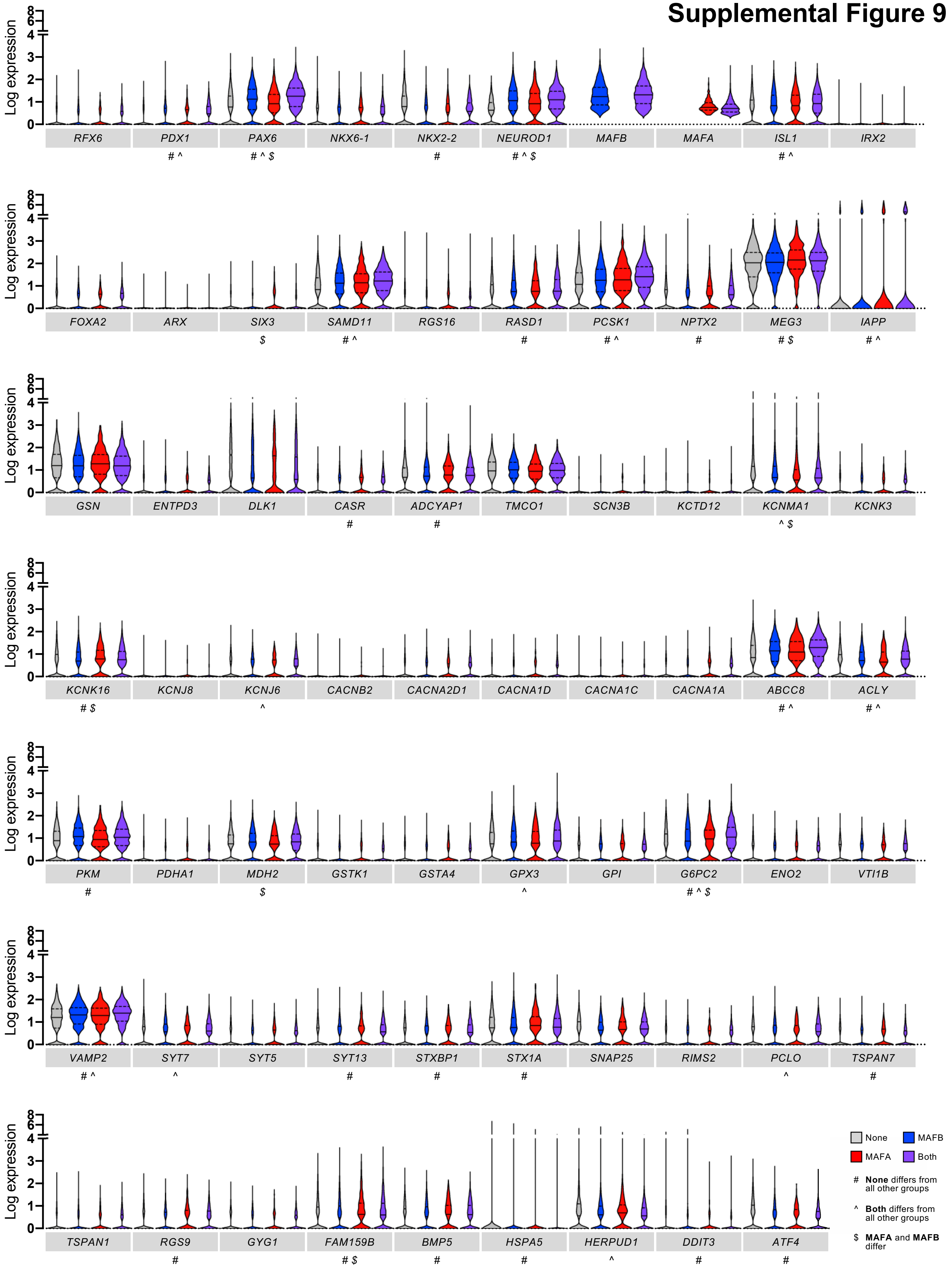

### Supplemental Figure 10

**A**

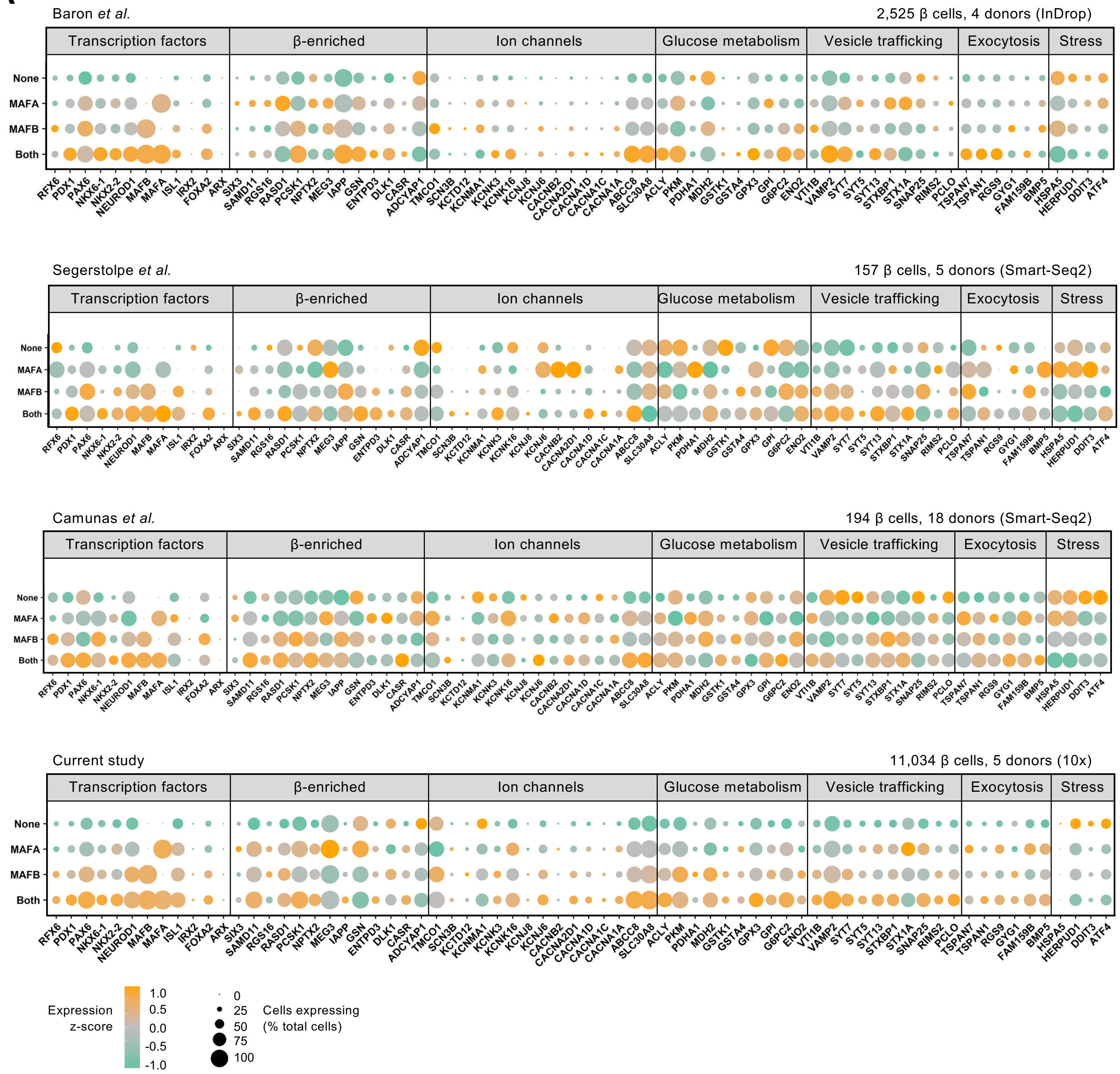


**B**

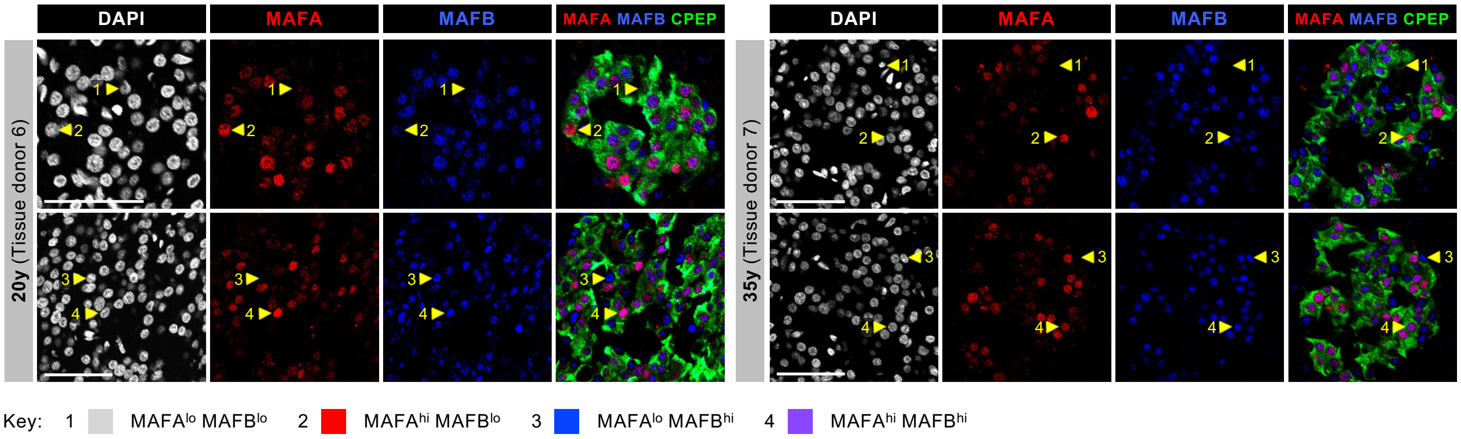

### Table S1. Human Islet Donor Information

**Checklist for Reporting Human Islet Preparations Used in Research**
Adapted from Hart NJ, Powers AC (2018) Progress, challenges, and suggestions for using human islets to understand islet biology and human diabetes. Diabetologia https://doi.org/10.1007/s00125-018-4772-2.

| **Islet preparation** | **1** | **2** | **3** | **4** | **5** |
|---|---|---|---|---|---|
| Unique identifier | DON184 | DON185 | RRID: SAMN08768781 | RRID: SAMN08768783 | R232 |
| Donor age (years) | 14 | 39 | 50 | 59 | 66 |
| Donor sex | Female | Female | Male | Female | Female |
| Donor BMI (kg/m$^2$) | 24.13 | 34.76 | 22.4 | 32.3 | 18.5 |
| Donor HbA1c (ng/mL) | 5.4 | 4.7 | N/A | N/A | 6.1 |
| Origin/source of islets | OPO | OPO | IIDP | IIDP | ADI |
| Islet isolation center | University of Pennsylvania | University of Pennsylvania | University of Pennsylvania | University of Wisconsin | University of Alberta |
| Donor history of diabetes? Yes/No | No | No | No | No | No |
| Donor cause of death | Anoxia/ Cardiovascular | Anoxia/ Drug intoxication | Cerebrovascular/ Stroke | Cerebrovascular/ Stroke | Cerebrovascular/ Stroke |
| Glucose-stimulated insulin secretion or other functional measurement | Perifusion | Perifusion | Perifusion | Perifusion | Perifusion |
| Handpicked to purity? Yes/No | Yes | Yes | Yes | Yes | Yes |
| Warm ischaemia time (hours) | N/A | N/A | No | No | N/A |
| Cold ischaemia time (hours) | 12 | 8.5 | 9.9 | N/A | N/A |
| Estimated purity (%) | 75 | 95 | 95 | 95 | 90 |
| Total culture time (hours) | 36 | 96 | 19 | 44 | 144 |

*ADI, Alberta Diabetes Institute; IIDP, Integrated Islet Distribution Program; N/A, not available;* OPO, *Organ Procurement Organization*

#### Table S2. Markers used for cell type annotation

| Cell type | Gene marker(s) |
|---|---|
| α cell | *GCG* |
| β cell | *INS* |
| δ cell | *SST* |
| γ cell | *PPY* |
| ε cell | *GHRL* |
| Acinar | *PRSS1* |
| Ductal | *KRT19* |
| Stellate | *PDGFRB, COL1A1* |
| Endothelial | *PECAM1* |
| Immune | *HLA-DRA* |

#### Table S3. Antibodies used for immunohistochemistry and flow cytometry

| Antigen/Conjugate | Species | Source | Catalog # | Application | Dilution |
|---|---|---|---|---|---|
| ARX | Sheep | R&D Systems | AF7068 | IHC (1º) | 1:2000 |
| C-peptide | Rat | Developmental Studies Hybridoma Bank | GN-ID4 | IHC (1º) | 1:200 |
| NTPDase3 (CD39L3) | Mouse | J. Sévigny | N/A | FC (1º) | 1:50 |
| Glucagon | Mouse | Abcam | ab10988 | IHC (1º) | 1:100 |
| HPa3 (HIC3-2D12) | Mouse | P. Streeter/M. Grompe | N/A | FC (1º) | 1:200 |
| HPi1 (HIC0-4F9) – biotin | Mouse | Novus | NBP1-18872B | FC (1º) | 1:100 |
| MAFA | Rabbit | Novus | NBP1-00121 | IHC (1º) | 1:250 |
| MAFB | Mouse | R&D Systems | MAB3810 | IHC (1º) | 1:1000 |
| Mouse Ig – APC | Goat | BD Biosciences | 550826 | FC (2º) | 1:500 |
| Mouse IgG – Cy3 | Donkey | Jackson Immunoresearch | 715-165-150 | IHC (2º) | 1:500 |
| Mouse IgG – Cy5 | Donkey | Jackson Immunoresearch | 715-175-150 | IHC (2º) | 1:300 |
| Mouse IgM – PE | Goat | Jackson Immunoresearch | 115-116-075 | FC (2º) | 1:1000 |
| Rabbit IgG – Cy3 | Donkey | Jackson Immunoresearch | 711-165-152 | IHC (2º) | 1:500 |
| Rabbit IgG – Cy5 | Donkey | Jackson Immunoresearch | 711-175-152 | IHC (2º) | 1:300 |
| Rat IgG – Cy2 | Donkey | Jackson Immunoresearch | 712-225-153 | IHC (2º) | 1:500 |
| Sheep IgG – Cy2 | Donkey | Jackson Immunoresearch | 713-225-147 | IHC (2º) | 1:500 |
| Streptavadin – BV420 | N/A | BD Biosciences | 563259 | FC (2º) | 1:500 |

*1º, primary antibody; 2º, secondary antibody; FC, flow cytometry; IHC, immunohistochemistry*

### Table S4. Human Pancreatic Tissue Donor Information

**Checklist for Reporting Human Islet Preparations Used in Research**
Adapted from Hart NJ, Powers AC (2018) Progress, challenges, and suggestions for using human islets to understand islet biology and human diabetes. Diabetologia https://doi.org/10.1007/s00125-018-4772-2.

| Pancreatic tissue | 6 | 7 | 8 |
|---|---|---|---|
| Unique identifier | DON54 | DON42 | DON61 |
| Donor age (years) | 20 | 35 | 55 |
| Donor sex | Male | Male | Male |
| Donor BMI (kg/m$^2$) | 19.442 | 26.852 | 35.565 |
| Donor HbA1c (ng/mL) | 5.6 | 5.1 | Not recorded |
| Origin/source of tissue | IIAM | IIAM | IIAM |
| Donor history of diabetes? Yes/No | No | No | No |
| Donor cause of death | Head trauma | Head trauma | Cerebrovascular/Stroke |
| Glucose-stimulated insulin secretion or other functional measurement | Not done | Not done | Not done |
| Warm ischemia time (hours) | DBD | DBD | No |
| Cold ischemia time (hours) | 6.2 | 15.1 | 26.6 |

*ADI, Alberta Diabetes Institute; DBD, Donation after Brain Death (minimal warm ischemia time); IIAM, International Institute for the Advancement of Medicine*
